# Strength of selection potentiates distinct adaptive responses in an evolution experiment with outcrossing yeast

**DOI:** 10.1101/2022.05.19.492575

**Authors:** Mark A. Phillips, Rupinderjit K. Briar, Marcus Scaffo, Shenghao Zhou, Megan Sandoval-Powers, Molly K. Burke

**Affiliations:** Department of Integrative Biology, Oregon State University Corvallis, OR 97331

**Author notes:** Corresponding authors: Mark A. Phillips, Molly K. Burke.

**Keywords:** experimental evolution, adaptation, genomics, population genomics, GxE

## Abstract

Experimental evolution studies with sexually-reproducing populations consistently find that adaptation is highly polygenic and fueled by standing genetic variation. However, studies vary substantially with respect to other evolutionary dynamics. Resolving these discrepancies is a crucial next step as we move toward extrapolating findings from laboratory systems to natural populations. Differences in experimental parameters between studies can perhaps answer these questions, and here we assess how one such parameter - selection intensity - influences outcomes. We subject populations of outcrossing *Saccharomyces cerevisiae* to zero, moderate, and high ethanol stress for ∼200 generations and ask: 1) does stronger selection lead to greater changes in allele frequencies at adaptive sites; and 2) do targets of selection vary with intensity? With respects to sites with large effects, we find some evidence for positive correlations between selection intensity and allele frequency change. While we observe shared genomic responses across treatments, we also identify treatment-specific responses. Combined with evidence of phenotypic trade-offs between treatments, our findings support the hypothesis that selection intensity influences evolutionary outcomes due to pleiotropic and epistatic interactions. We conclude that it should be a major consideration when attempting to generalize inferences across studies; in other words, we argue that different intensities of selection effectively create distinct environments and genotype-by-environment interactions. Lastly, our results demonstrate the importance of clearly-defined controls in experimental evolution. Despite working with a presumably lab-adapted model system, without this element we would not have been able to distinguish genomic responses to ethanol stress from those associated with laboratory conditions.

## Introduction

Evolve and resequence (“E&R”) experiments are now commonly used to study the genetics of adaptation and complex traits (Long et al. 2015; Schlötterer et al. 2015). In these studies, experimental populations are subjected to carefully designed selective regimes under controlled conditions, and sampled for DNA or RNA sequencing over many generations. This experimental framework allows for powerful statistical associations between observed, presumably adaptive, phenotypic responses with shifts in genetic variation or changes in gene expression. In addition to linking genotypes to phenotypes, the ability to control population size, level of starting genetic variation, and environmental conditions make these experiments a powerful tool for studying broad adaptive dynamics. It is now well understood that adaptation in E&R studies featuring outcrossing systems is fueled by standing genetic variation (Long et al. 2015; Barghi et al. 2020). However, the specific evolutionary dynamics observed across these studies vary considerably. We believe achieving a better understanding of what causes these differences is a key step towards extending general findings from E&R to real populations (Phillips and Burke 2021).

Major points of discrepancy in the literature today include topics like evolutionary repeatability, the role of contingency in the genetics of adaptation, and what some term “sweeps versus shifts” (Barghi et al. 2020). Conflicting findings around evolutionary repeatability in particular have received a great deal of attention. While some studies report high levels of evolutionary repeatability, as evidenced by parallel responses to selection across replicate populations (e.g. Linder et al. 2022), others find more idiosyncratic and replicate-specific responses (e.g. Barghi et al. 2019). While this issue might not be entirely resolved, comparing across studies does point to likely explanations. As synthesized by Schlötterer et al. (2023), differences in trait architecture (i.e. simple versus complex), number of founder genotypes, and properties of founder populations (e.g. levels of linkage disequilibrium and polymorphism) are likely driving observed differences in evolutionary repeatability. More broadly, we believe this example of repeatability highlights the need to better understand ways various experimental parameters shape evolutionary outcomes before extending inferences from E&R studies to populations outside the lab. Here we focus on how an often overlooked parameter – selection intensity – impacts evolutionary outcomes in large outcrossing populations with abundant standing genetic variation.

How a given selective pressure manifests (e.g. intensity, direction, duration, constant versus dynamic, etc.) has clear ecological relevance that likely shapes evolutionary outcomes in both natural populations and laboratory experiments. While this topic has been explored across a number of theoretical studies (e.g. Kessner and Novembre 2015; Stetter et al. 2018; Christodoulaki et al. 2019; Vlachos and Kofler 2019; Hayward and Sella 2021), few E&R studies have sought to address it empirically. Recent work suggests the relationship between how selection is imposed and how populations respond is more complex than might have previously been expected. For example, Otte et al. (2021) compares responses to selection in cold and heat- stressed populations of *D. simulans*. As these are both forms of temperature stress, a simple expectation would be that targets of selection are common to both regimes but allele frequencies move in opposite directions in each. However, while they observed a similar number of targets under both selection regimes, overlap was limited and targets with the highest effect sizes were treatment-specific (implying that heat and cold resistance have distinct genetic bases). Here we aim to expand this body of work by testing whether or not varying the intensity of selection pressure in a single direction might similarly impact outcomes.

Using populations of outcrossing *S. cerevisiae* derived from a single ancestral population,, we compare the genomic response to moderate and high ethanol exposure. In the simplest scenario, one might expect a set of “ethanol resistance” alleles segregating in our ancestral population, which should increase in frequency in both selection treatments in proportion to selection intensity. Theoretical work supports this idea with findings from Christodoulaki et al. (2019) indicating that when new trait optima are distant (i.e. when selection is high-intensity), the genomic response involves larger sweep-like changes in and around targets of selection compared to more modest shifts when optima are close. However, if the genetics of complex traits is characterized by widespread pleiotropy as some suggest (Visscher and Yang 2016; Boyle et al. 2017) or epistasis, this might not be the case. For example, due to pleiotropic effects and resulting trade-offs with other characters, alleles that are adaptive under high ethanol stress conditions might not be favored at lower levels of exposure and vice-versa. Lastly, past work with outcrossing *S. cerevisiae* populations identified an entire class of alleles beneficial at late but not early stages of adaptation, providing some support for the idea that epistatic interactions may be common in this system (Phillips et al. 2020). If generally true, this could also drive treatment-specific responses - alleles beneficial at moderate levels of ethanol stress may not confer the same benefits at high levels of ethanol exposure due to negative interactions with alleles unambiguously beneficial under those conditions. These examples invoke different types of genotype-by-environment (hereafter, GxE) interactions that result from different selection intensities effectively creating distinct environments.

In summary, with this work we ask the following questions: (i) does more intense selection lead to larger changes in frequencies at and around target sites, and (ii) do targets of selection vary with selection intensity? We address these questions using genomic data from outcrossing *S. cerevisiae* populations subjected to zero, moderate, and high ethanol stress for ∼200 generations. We believe that a deeper understanding of how selection intensity influences trait architecture will enable not only more productive synthesis across E&R work, but also better-informed models of polygenic adaptation and an improved capacity to extrapolate results from specific studies to the real world.

## Materials and Methods

### Experimental populations and selection regimes

All experimental populations used in this study were derived from the “S12” population described in detail in Phillips et al. (2021). Briefly, this population was created by combining 12 haploid strains from the barcoded SGRP yeast strain collection (Cubillos et al. 2009). The specific strains used are: Y12, YSP128, SK1, DBVPG6044, UWOPS05_217_3, L_1528, L_1374, DBVPG6765, YJM981, YJM975, BC187, and N273614. As described in Linder et al. (2020), these strains have been modified to enable easy crossing and diploid recovery – these modified strains were kindly provided by Dr. Anthony Long. The MATa and MATα strains contain *ho* deletions to prevent mating-type switching, but each contain a different drug-resistance marker inserted into a pseudogene (YCR043C) tightly linked to the mating-type locus (*MATa, ho*Δ*, ura3::KanMX-barcode, ycr043C::NatMX* and *MAT*α*, ho*Δ*, ura3::KanMX-barcode, ycr043C::hphMX*). These genotypes enable haploids of each mating type to be recovered using media supplemented with either hygromycin B or nourseothricin sulfate, and they enable newly mated a/α diploids to be recovered in media supplemented with both drugs.

To create the S12 recombinant outbred population from these haploid strains, we used a semi-round-robin design of initial crossing, followed by 12 iterations of what we call our “weekly outcrossing protocol” to maximize genetic diversity and allow for some domestication to laboratory handling conditions (detailed in Burke et al. 2020; Phillips et al. 2021). Due to the already high species-specific recombination rates (Liu et al. 2019), as well as this extra selection for high rates of outcrossing, regions of linked alleles that may potentially respond to experimental evolution are expected to be relatively small in the population (as small as 5-10KB; *cf.* Burke 2023). The experimental populations featured in this study were then derived from samples of this base population: 20 control replicates (C_1-20_), 20 moderate ethanol stress replicates (M_1-20_), and 20 high ethanol stress replicates (H_1-20_). Levels of ethanol used in the two stress treatments were chosen based on the results of exploratory growth rate assays (see next section for general assay methods). We found that 10% ethanol was close to the limit of what could support population growth (at least for a 48-hour period), and we thus chose this as the high stress treatment. We chose the moderate treatment to include 6% ethanol as this resulted in a doubling time approximately mid-way between the high stress and control conditions (Supplementary Figure 1, note: this plot is based on assays performed prior to the start of the experiment).

The weekly outcrossing protocol described by Burke et al. (2020) was also used to maintain sexual reproduction in these 60 replicate populations, with minor modifications to increase throughput (i.e. smaller volumes). This protocol involves batch culture of diploids in 1mL of liquid medium in alternating wells of 24-well plates (Corning); every other well contained sterile YPD which was monitored for growth (which would indicate potential cross-contamination) throughout the experiment. After batch culture of diploids, the entire 1mL of culture was washed and resuspended in 1mL minimal sporulation media (1% potassium acetate), and incubated with shaking for 72 hours (30°C/200 rpm). After sporulation, a modified random spore isolation protocol was implemented to disrupt asci and isolate spores. This protocol involves resuspending sporulated cultures in 1mL Y-PER Yeast Protein Extraction Reagent (Thermo), followed by incubation at 50°C for 15 minutes to kill vegetative diploid cells.

Cultures were then resuspended in a 1% zymolyase (Zymo Research) solution to weaken ascus walls, and vortexed at maximum speed with 0.5 mm silica beads (BioSpec) to mechanically agitate the asci. Following these steps, spores were transferred to YPD agar plates supplemented with nourseothricin sulfate (100mg/L), hygromycin B (300mg/L) a nd G418 (200mg/L) and incubated at 30°C for 48 hours, during which time spores mated (due to close proximity on the plate) and diploids germinated. The resulting lawns of new diploid cells were scraped off plates using sterile glass slides and transferred to 10mL of sterile YPD media (1/10^th^ of this culture was preserved in 15% glycerol and archived at -80°C for DNA extraction and sequencing). From these super-saturated cultures, 10 μL were transferred into 1mL of treatment-specific medium and incubated for 48 hours (30C/200rpm), with a 1/100 dilution halfway through to increase generational turnover. For the control treatment, standard YPD (2% yeast extract, 1% peptone, 2% dextrose) was used as the culture medium. For the moderate and high ethanol stress treatments, YPD was supplemented with either 6% or 10% ethanol (by volume), respectively. After this period of batch culture, the next weekly outcrossing iteration would commence (Supplementary Figure 2). All replicates were handled in parallel for a total of 15 weeks, with one cycle of outcrossing per week.

By estimating cell density at various steps of the outcrossing protocol (via OD_600_ of liquid cultures, and/or colony counts of dilution series), we can approximate the number of asexual generations that occurred in each treatment during the experiment. We estimate that ∼10 cell doublings occurred during the 48-hour period of batch culture – this phase of competitive growth in treatment-specific medium is most relevant to our major questions relating to selection strength. We infer that an additional ∼4 cell doublings occur during the non-competitive phases of the protocol (i.e. growth on agar medium). We observed that growth was slower (and cell densities lower) in the high ethanol stress treatment compared to the others, but this did not lead to dramatically different estimates of cell doublings among the treatments. Ultimately, we project that over 200 asexual generations elapsed in all treatments over 15 weeks. This generational turnover is difficult to precisely estimate for several reasons, chiefly because some aspects of population biology were not routinely quantified throughout the experiment. For example, our protocol development (*cf*. Burke et al. 2020) suggests that sporulation and mating efficiencies are high in the base population, and that the mortality induced by the weekly outcrossing protocol is low. But it was not possible to quantify these metrics in all 60 experimental replicates every week. If sporulation efficiency, mating efficiency, and/or spore viability was low in a particular replicate during a given week, this would reduce the number of viable cells in the population (compared to the baseline expectation) and lead to a higher estimate of cell doublings for the week. Thus, we feel that our ballpark estimates can be considered rather conservative.

### Growth rate assays

Growth in liquid medium was the primary phenotype we tracked in this experiment. We compared growth between evolved populations and the ancestor in each media type to detect evidence of phenotypic adaptation in a given treatment. We also compared growth of evolved populations across all media types to detect evidence of potential phenotypic trade-offs.

For each population assayed, a culture would first be grown overnight in 10 mL YPD in a shaking incubator (30 °C/200 rpm). Cultures were then quantified via absorbance at OD_600_ (values of 1/100 dilutions typically ranged from 0.10 to 0.15 across all treatments and timepoints) and diluted to a standard starting concentration of OD_600_ = 0.1 using the appropriate media type. These dilutions were then transferred to individual wells of a 96-well plate (Corning) in 200 μL volumes. A Tecan Spark multimodal microplate reader measured absorbance in each well every 30 minutes over a 48-hour period at 30°C with no shaking. Measurements were taken at four positions within each well and the average values were used for subsequent data analysis. To avoid edge effects, wells on the outer perimeter of the plates were not used for assays and simply filled with sterile YPD. In a given plate reader assay, we included 9 technical replicates of the ancestral population and 10 randomly selected replicates from each treatment. Using these protocols, independent plate reader assays were run under control, moderate ethanol stress, and high ethanol stress conditions.

Using the data from our plate reader runs and 7he R (R CoreTeam 2021) package “Growthcurver” (Sprouffske and Wagner 2016) we estimated doubling times and carrying capacity. Estimates were obtained by fitting data to the following logistic equation that gives the number of cells *N_t_* (as measured by absorbance) at time *t*:

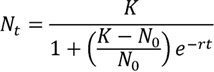

Where starting population size is represented by *N_0_*and carrying capacity by *K*. Here carrying capacity is simply defined as the maximum population size in a particular environment. Lastly, *r* represents the growth rate that would occur if there were no limits on total population size. This value is also used to calculate the doubling time which is defined as 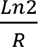. From here, statistical comparisons were done on a per-assay basis. For a given assay, we used Kruskal-Wallis tests to compare mean carrying capacities and doubling times, and pairwise Wilcoxon rank sum tests to determine which specific groups were significantly different from one another. We chose to carry out our analysis in this manner to avoid the confounding effects of run-to-run variation.

While qualitative patterns are consistent between runs, estimates like carrying capacity can vary a great deal between runs under the same conditions.

### DNA extraction, sequencing, and read mapping

The 20 replicates of each treatment were sampled for pooled-population genome sequencing at three distinct timepoints: after a single outcrossing cycle, after 7 outcrossing cycles, and after 15 outcrossing cycles. To sample populations for sequencing, 1 mL of freezer stocks were revived on YPD agar plates. After 48 hours of growth at 30°C, the resulting lawns were broadly sampled by wooden applicator (to capture as much genetic diversity as possible) and these cells were inoculated into 10mL of liquid YPD culture, and grown overnight in the shaking incubator. Genomic DNA was extracted from samples with Qiagen’s Yeast/Bact Kit following the manufacterer’s protocol. After checking DNA quantity, sequencing libraries were prepared for Illumina sequencing with the Nextera Kit DNA Sample Preparation Kit, implementing some routine modifications to increase throughput (e.g. Baym et al. 2015).

Libraries were pooled into groups of 48 and run on at least one PE150 lane of the HiSeq3000 housed at OSU’s Center for Quantitative Life Sciences (CQLS); samples with lower-than-average coverages were re-quantified, re-pooled and re-sequenced such that high (>50X average genome-wide) coverages were achieved across all experimental replicates.

The SNP calling pipeline we routinely use has been described previously (e.g. Phillips et al. 2020. We used GATK v4.0 (Van der Auwera & O’Connor 2020) to align raw data to the *S cerevisiae* S288C reference genome (R64-2-1) and create a single VCF file for all variants identified across all populations, using standard best practices workflows and default filtering options. This VCF file was converted into a “raw” SNP frequency table by extracting the AD (allele depth) and DP (unfiltered depth) fields for all SNPs passing quality filters; the former field was used as the alternate allele count observed at a presumed SNP, and the latter was used as the total coverage observed at that site. The VCF file was also used as an input for SnpEff v4.3 (Cingolani et al. 2012) to extract potential functional effects of individual SNPs. SnpEff annotates each variant in a VCF file (e.g. by tagging whether it occurs within a protein-coding sequence) and calculates the effect(s) each produces on known genes (e.g. amino acid changes).

We applied several filtering and quality control steps to the raw SNP table generated from the steps above to ensure only high-confidence sites were used in subsequent analysis. Mean genome-wide coverage across the 180 samples sequenced for this study ranged from ∼50X to ∼300X with a median value of ∼80X (Supplementary Table 1). We began our filtering process by only considering sites where sequence coverage exceeded 20X in all populations. Next, we removed all sites that were not expected to be polymorphic based on the previously described sequences of the founder strains used to create the ancestral population (*cf.* Phillips et al. 2021). We also removed sites that were not polymorphic in the ancestral population itself, as for this study our objective is to track the evolution of standing genetic variants, and not *de novo* mutations. Finally, we only considered sites where the alternate nucleotide frequency fell between 0.02 and 0.98 across the entire dataset. This removes sites where the alternate nucleotide is fixed across all populations, and sites where errors in sequencing and/or variant calling may create the appearance of polymorphism. These steps ultimately resulted in a SNP table with 61,281 high-quality SNPs.

### Identifying candidate sites and selection response categories

In pursuing the goal of associating particular candidate SNPs with adaptation, we found it useful to assign these to one of five distinct categories: (i) general adaptation to laboratory handling, (ii) general adaptation to ethanol stress, (iii) specific adaptation to the high ethanol treatment, (iv) specific adaptation to the moderate ethanol treatment, and (v) specific adaptation to the control treatment. Comparing the magnitude of change at sites across these categories allows us to assess whether greater selection intensity results in greater shifts in allele frequencies, and comparing the location of sites observed across these categories allows us to assess whether or not different selection intensities involve different genomic targets. Notably, the decision to include category (v) above was made *after* observing evidence of continued and unique adaptation in our control treatment; this was an unanticipated outcome that has thematically influenced our interpretation of all other results. To identify and differentiate between these types of responses we used a combination of within- and between-treatment comparisons. For within-treatment comparisons (e.g. comparing SNP frequencies between cycle 1 and cycle 15 for the C_1-20_ populations), we relied on the Cochran-Mantel-Haenszel (CMH) tests. For between-treatment comparisons (e.g. comparing SNP frequencies between C_1-20_ and H_1-20_), we used a generalized linear mixed model (GLMM) approach.

Patterns of overlap between these different comparisons were then used to isolate the different types of responses we were interested in (see Table 1 for details on how we define each type of response). As coverage variation can impact statistical results when using nucleotide counts, as both these methods do, we scaled coverage across the genome to a uniform level of 50X before performing these tests (Wiberg et al. 2017). Importantly, we chose approaches best-suited to the detection of parallel responses to selection across replicates, as we believe such sites are most likely to be true targets of selection.

**Table 1.** The five major response types used to categorize candidate sites. These are defined based on how sites significant in tests of allele frequency differentiation overlap within and between treatment-specific statistical comparisons.

| Response type | Definition | Number of candidate SNPs |
| --- | --- | --- |
| General laboratory selection | Significantly differentiated sites that overlap across all three treatments based on comparisons between cycles 1 and 15 for each. Sites differentiated between C vs H and M treatments at cycle 15 were removed to account for directionality. | 695 |
| General ethanol selection | Significantly differentiated sites that overlap across H and M treatments based on comparisons between cycles 1 and 15 for each. Any sites also significant in the C treatment were removed, along with any sites differentiated between the H vs M treatment at cycle 15. | 605 |
| High ethanol-specific | Significantly differentiated sites unique to the H treatment based on cycle 1 vs cycle 15 comparisons (i.e. sites significant in other cycle 1 vs 15 comparisons were removed). | 477 |
| Moderate ethanol-specific | Significantly differentiated sites unique to the M treatment based on cycle 1 versus cycle 15 comparisons (i.e. sites significant in other cycle 1 vs 15 comparisons were removed). | 591 |
| Control-specific | Significantly differentiated sites unique to the C treatment based on cycle 1 versus cycle 15 comparisons (i.e. sites significant in other cycle 1 vs 15 comparisons were removed). | 857 |

The CMH tests for within-treatment comparisons are commonly used in E&R studies, and simulations suggest this is one of the most accurate methods available for identifying responses to selection based on sampling populations over time (Vlachos et al. 2019). Tests were performed using the “poolSeq” package (Taus et al. 2017) in R. For a given within-treatment comparison, the data were paired appropriately (e.g. C_1_ cycle 1 with C_1_ cycle 15, C_2_ cycle 1 with C_2_ cycle 15, and so on for the control treatment) and tests were performed at all 61,281 K SNPs. Significant sites show a consistent change in frequency from cycle 1 to cycle 15, across all replicate populations. As we sequenced populations at an intermediate timepoint (cycle 7) we also had the option to use more sophisticated time-series approaches for this task. However, given that simulations (Vlachos et al. 2019) and empirical comparisons (Phillips et al. 2020) have shown that these approaches perform similarly to the CMH test using starting and ending points in an E&R study, we favored the latter approach.

For between-treatment comparisons, a core assumption of the CMH test is violated as there is no meaningful way to pair replicate populations (e.g. pairing C_1_ with H_1_ is no more meaningful than C_1_ with H_2_). As such, the CMH test is not appropriate and we instead opted to use the GLMM approach, which emphasizes parallel changes across replicate populations, and is becoming increasingly popular in E&R studies (e.g. Jha et al., 2015, Kawecki et al., 2021). Three independent pairwise comparisons were performed using this approach: C_1-20_ versus H_1-20_, C_1-20_ versus M_1-20_, and H_1-20_ versus M_1-20_ (note: these comparisons only use cycle 15 data). For a given comparison, the “lme4” package (Bates et al. 2015) was used to fit a GLMM with binomial error structure to nucleotide counts at a given polymorphic site. The following model statement was used:

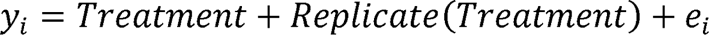

where y_l_ is the frequency of the *i*th SNP, Treatment is a fixed effect, and Replicate is a random factor nested within Treatment, and e_l_ refers to the binomial error term. To test for significance, the “anova” function in R was used to compare the full model to a simplified model without the fixed effect. Using this approach, we were able to identify SNPs that were consistently differentiated between treatments.

To correct for multiple comparisons and establish significance thresholds, we used the same general permutation approach described in Burke et al. (2014) and Graves et al. (2017). Using C_1-20_ cycle 1 versus C_1-20_ cycle 15 as an example: 1) samples were randomly assigned to one of two groups, 2) the CMH tests were performed at each SNP in the shuffled data set, and 3) the smallest p-value generated was recorded. This was done 1000 times, and the “quantile” function in R was used to determine the cutoff for the 0.5th percentile. This was then used as the significance threshold for this comparison; that is to say, a Type I error rate of 0.005 is required for a site to be considered significantly differentiated. Here we use a 0.005 significance threshold instead of the typical 0.05 as this increased stringency has been suggested to improve reproducibility (Benjamin et al. 2018). This process was repeated with the M and H populations to establish significance thresholds for the M_1-20_ cycle 1 versus M_1-20_ cycle 15 and H_1-20_ cycle 1 versus H_1-20_ cycle 15 CMH comparisons. The same general procedure was also used for between-treatment comparisons using the GLMM approach. However, for GLMM comparisons, we only ran 200 permutations instead of 1000 as this method is more computationally intensive than the CMH. We did perform a trial using 1000 permutations for one comparison and found that it produced a very similar threshold.

In principle, response categories could have been defined only using the CMH results.

However, incorporating results from between-treatment comparisons allowed us to further refine general ethanol and laboratory selection categories. To give an example, general ethanol selection candidates would include significant sites that overlap between the M and H within treatment comparisons. However, it is possible this could include sites that are actually moving in opposite directions in these treatments. Removing sites that are differentiated between H and M therefore allows us to more strictly define general ethanol selection candidates as those that show a shared directional response in both treatments.

### Estimating selection coefficients

In an effort to further characterize genomic responses to selection, we estimated selection coefficients (*s*) across the genome in each treatment based on changes in SNP frequencies over time. These were generated using Bait-ER (https://github.com/mrborges23/Bait-ER). Bait-ER is a Bayesian approach to estimate selection coefficients and identify targets of selection from E&R timeseries data developed by Barata et al. (2023). We chose Bait-ER specifically as findings from Barata et al. (2023) suggest *s* estimates are accurate even when effective population size (N_e_) is miss-specified by orders of magnitude. We believe this is an important consideration for our study as populations feature complex life cycles which violates many of the assumptions underlying standard methods for estimating N_e_ from E&R timeseries data (e.g. Jónás et al. 2016). For instance, our populations experience sexual and asexual generations, census size can vary greatly between these phases, and our focal selective pressure is only present during the generations in batch culture.

Using Bait-ER we estimated *s* based on changes in SNP frequencies over all timepoints sampled for each treatment. However, we first needed to estimate N_e_ for each group. This was done using the method described by Jónás et al. (2016) and implemented in the “poolSeq” package in R developed by Taus et al. 2017. Specifically, estimates were generated using SNP frequencies from the first and final timepoint sampled for each population using the “estimateNe” function with “method = P.planII”. This method was designed specifically to account for the two-stage sampling process associated with pool-seq data like ours. We specified that the number of generations between timepoints (t) was 200, and number of individuals sampled (poolsize) was 1*10^6^ (note: as this poolsize is only a rough estimate, we tried a range of values and found that differences in either direction by several orders of magnitude did not affect estimates). Finally, we averaged N_e_ estimates across replicate populations to get a single estimate for each treatment (Supplementary Table 2) that was then used for Bait-ER to generate *s* estimates for polymorphic sites in each group. Ranging from 350 in the C populations to 470 in the H populations, we do find that N_e_ are far below what we would expect and attribute this to the factors described above. As such, we only interpret N_e_ estimates in a relative sense.

### Selection intensity and magnitude of allele frequency change

To assess the prediction that more intense selection should result in greater changes in allele frequencies at target sites, we simply compared the mean changes in SNP frequency at candidate sites between treatments. If this idea is correct, among sites associated with ethanol resistance, we would expect to see significantly greater changes in SNP frequency in the high ethanol stress treatment compared to the moderate ethanol stress treatment. This should extend to both the specific targets of selection and linked sites.

### Gene ontology (GO) term analysis

Once SNPs belonging to each category of selection response listed in Table 1 were identified, we compared these for potential differential enrichment of gene ontology (GO) terms. These lists were created using the SnpEff output: for each response type, we created a list of genes associated with at least one candidate SNP in that category (note: here we consider all effect types as valid associations). Text files with candidate list and associated SnpEff annotation are available through Dryad for interested individuals (see Data Availability statement for details). GO term enrichment analyses for each gene list were then performed using Metascape (Zhou et al. 2019) with default settings. For our analysis, a gene was only counted once, even if there were multiple candidate SNPs associated with it. Our analysis was agnostic to which SNPs were the true target of selection and multiple candidates associated with a gene were assumed to be a result of linkage.

## Results

### Adaptation and growth rates

To characterize the long-term consequences of each experimental treatment, we independently assayed the growth of evolved (cycle 15) replicates in all three media types (control, moderate ethanol, high ethanol) and compared these to the growth of the ancestral population (Figure 1). To assess statistical differences, we used Kruskal-Wallis tests to compare mean values across all groups for a given assay, and pairwise Wilcoxon rank sum tests to determine which treatment pairs were significantly different (note: the Benjamini-Hochberg procedure was used to correct for multiple comparisons). We find that in plain YPD, differences in doubling time and carrying capacity are small (Figure 1A and B, Supplementary Figure 3A), but significant for both doubling time (Kruskal-Wallis test p=0.0001) and carrying capacity (p= 0.001). In the case of doubling time in YPD, there are no significant differences between the C, M, and H replicates and the variation between biological replicates is small. However, the ancestral population has a significantly longer doubling time than evolved populations (mean of 1.3 hours for the ancestor versus 1.25 to 1.26 for C, M, and H populations); this implies that replicates in all treatments adapted to laboratory conditions in ways that improved growth in YPD. For carrying capacity in YPD, the H replicates appear to stop doubling at a lower cell concentration compared to the ancestral population, and the C and M treatments (mean of 0.9 in the H replicates versus ∼1 for the others, note: carrying capacity is represented as log(OD) as shown in Supplementary Figure 3). While we do find some statistically significant differences here, they are clearly quite small. As such, it appears that adaptation to ethanol stress in the moderate and high ethanol stress has not had a large negative impact on the ability of replicates in these groups to grow in YPD (i.e. no obvious suggestion of an adaptive trade-off).

**Figure 1.**
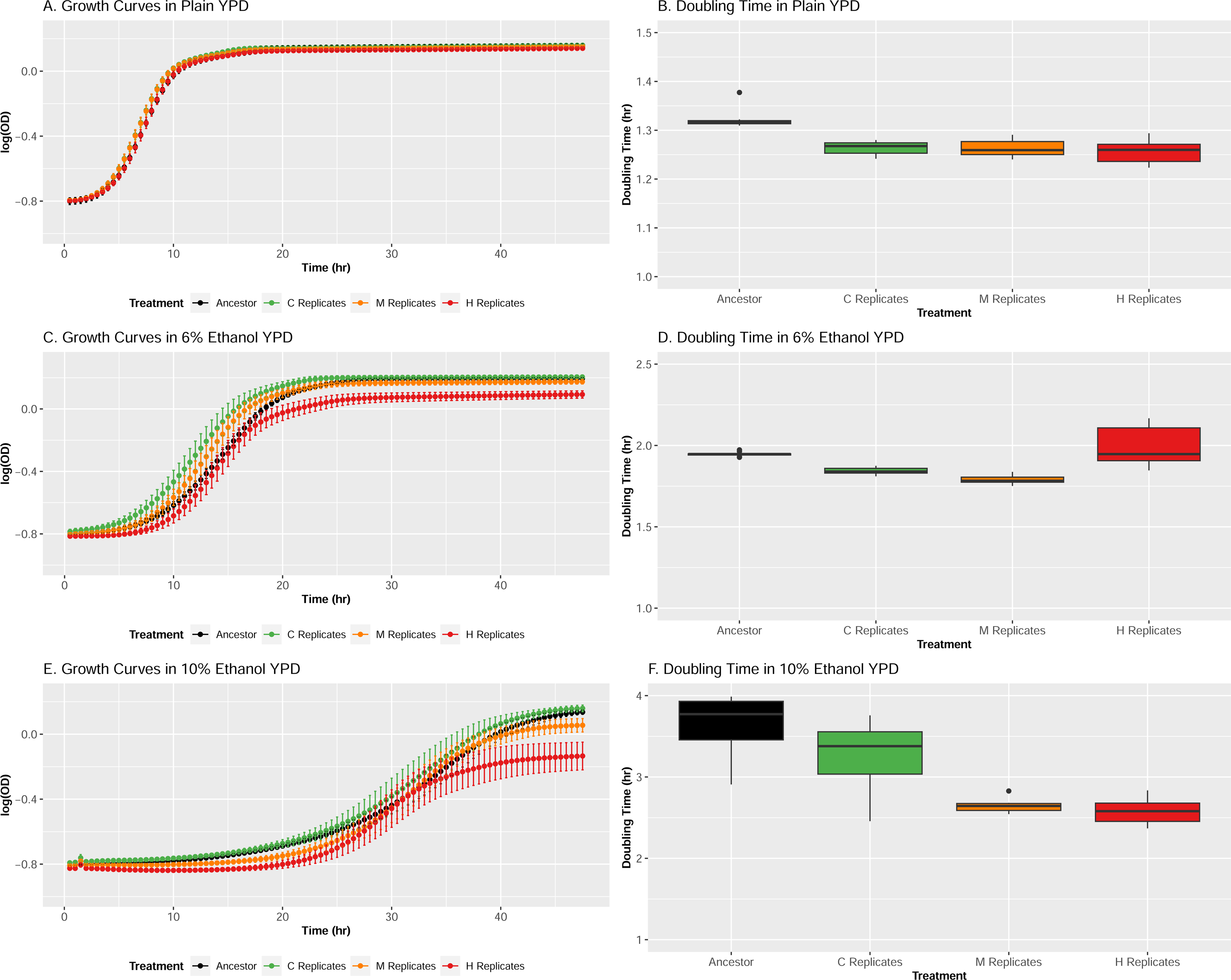
***Phenotypes of evolved populations.*** Growth rates and doubling times for the ancestral and experimental populations after 15 cycles of adaptation in control (A and B), moderate ethanol (C and D), and high ethanol conditions (E and F). In each assay, nine technical replicates of the ancestor and 10 randomly-chosen replicates from each treatment were evaluated.

We do observe differences in growth patterns among treatments in 6% ethanol (Figure 1C), with respect to both doubling time (Figure 1D, Kruskal-Wallis test p =1.75x10^-6^) and carrying capacity (Supplementary Figure 3B, Kruskal-Wallis test p=8.05x10^-7^). Both the M and C treatments grow significantly faster than the ancestral population, evidencing adaptation. By contrast, doubling time in the H treatment could not be significantly distinguished from the ancestor, and was correspondingly slower than the C and M treatments. Carrying capacity is significantly different in all pairwise comparisons except for the ancestor vs. C treatment.

Carrying capacity is lowest in the H treatment (mean=0.98) followed by the M treatment (mean = 1.07), and the C treatment (mean=1.1) resembled that of the ancestor (mean=1.1).

In populations assayed in 10% ethanol (Figure 1E), we also find significant differences in both doubling time (Figure 1F, Kruskal-Wallis test p=1.16x10^-5^) and carrying capacity (Supplementary Figure 3C, Kruskal-Wallis test p=5.46x10^-7^) among treatments. With respect to doubling time, we find significant differences between all pairwise comparisons except for the M replicates versus the H replicates. Doubling time is slowest in the ancestral population (mean = 3.64 hours), followed by the C replicates (mean = 3.26 hours), followed by the M and H replicates (mean = 2.64 and 2.57 hours). For carrying capacity, all pairwise comparisons are significantly different except for the ancestral population versus the C replicates. The ancestral population has the highest carrying capacity (mean = 1.54, same for the C replicates), followed by the M replicates (1.11) and H replicates (mean = 0.78). So, while populations adapted to ethanol stress treatments generally have faster growth rates under high ethanol stress conditions, they also stop doubling sooner. And adaptation to high ethanol stress specifically is associated with the lowest carrying capacity.

While these growth rate assays are useful for ranking phenotypic differences among ancestral and evolved populations (and they are internally consistent with respect to a given plate reader assay), we do not believe the raw metrics reported necessarily reflect what occurred during the actual selection experiment. For instance, in the plate reader assays, populations in 10% ethanol appear to still be in lag phase after nearly 24 hours of growth (Figure 1E). While we observed that replicates of the H treatment grew more slowly than the other treatments throughout the experiment, routine OD_600_ readings indicate that all populations were in log phase at 24 hours. So, whenever cultures were transferred to fresh liquid media during the experiment, cells were in log phase in all treatments; we felt that it was important to avoid sampling cells at different stages as this could introduce unwanted variation among treatments. There are many reasons why the absolute values of growth parameters estimated during the phenotype assays might differ from what was observed in the selection experiment, but we suspect the main drivers are volume differences (200 μL in 96 well plates versus 1mL in 24 well plates), and agitation (no shaking in the plate reader versus 200 rpm in the shaking incubator).

### Evidence for different response categories among candidate SNPs

To assess the genomic response to selection, we first compared SNP frequencies between cycles 1 and 15 for each treatment using the CMH test (Supplementary Figure 4). Here we find clear responses to selection as evidenced by consistent changes in SNP frequencies in each treatment, including the controls. As the number of candidate regions overlap across both ethanol stress treatments and controls, it appears that adaptation to general laboratory handling (i.e., our weekly outcrossing protocol) is a major feature of this study. To better parse these responses to selection, we used a GLMM approach to identify differentiated SNPs between treatments during cycle 15 (Supplementary Figure 5). Based on patterns of overlap between all of these comparisons, we identified 5 major response types: (i) general adaptation to laboratory handling, (ii) general adaptation to ethanol stress, (iii) specific adaptation to the high ethanol treatment, (iv) specific adaptation to the moderate ethanol treatment, and (v) specific adaptation to the control treatment. (Table 1). As seen in Figure 2 where these categories are overlaid on the CMH results comparing cycle 1 and 15 for each treatment, almost all significant SNPs can be assigned to one of these 5 categories.

**Figure 2.**
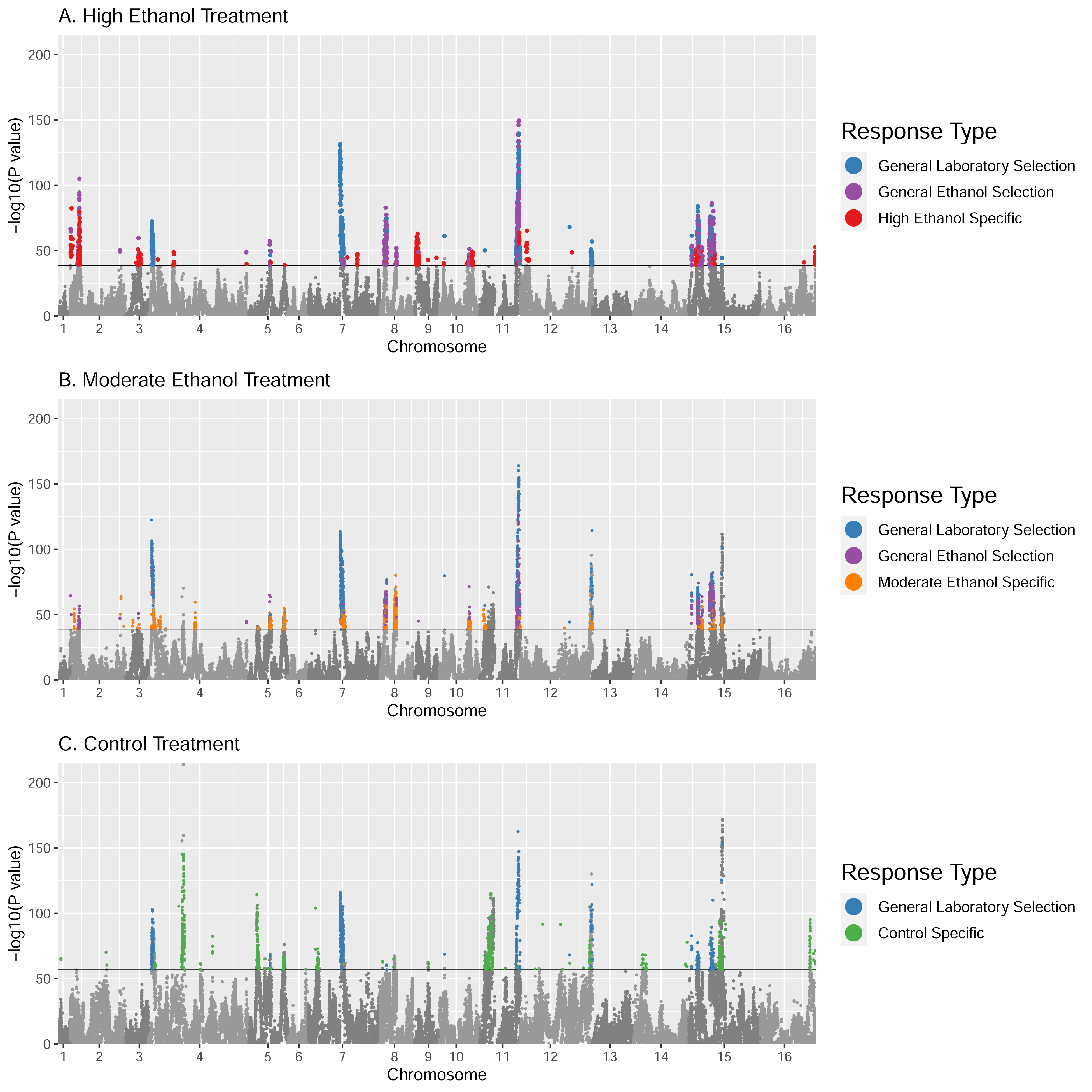
***Treatment-specific candidate regions.*** Results from CMH tests comparing SNP frequencies between cycles 1 and 15 in replicate populations selected for (A) high ethanol stress, (B) moderate ethanol stress, and (C) control conditions. Black lines represent permutation-derived significance thresholds, and color coding of significantly differentiated sites corresponds to the response categories described in Table 1.

Considering the major peaks in Figure 2, we do observe some mixing of categories within certain peaks, but most are either made up of a single response type or have a clear majority type. And while there are a number of regions that correspond to our three ethanol response types (general, high, and moderate ethanol responses), our strongest signals are observed in regions associated with general laboratory selection. So, while ethanol exposure clearly imposes some stress, laboratory conditions appear to be major selective pressures on their own. Here it should be noted that the ancestral population sampled to establish all experimental replicates had already experienced 12 “domestication” cycles with outcrossing prior to this experiment. As such, there was some expectation that these populations would already be adapted to routine culture and outcrossing protocols. A principal component analysis (PCA) of SNP frequencies indicates that despite what appears to be the shared selection response imposed by laboratory conditions, groups are still broadly differentiated by cycle 15 (Figure 3).

**Figure 3.**
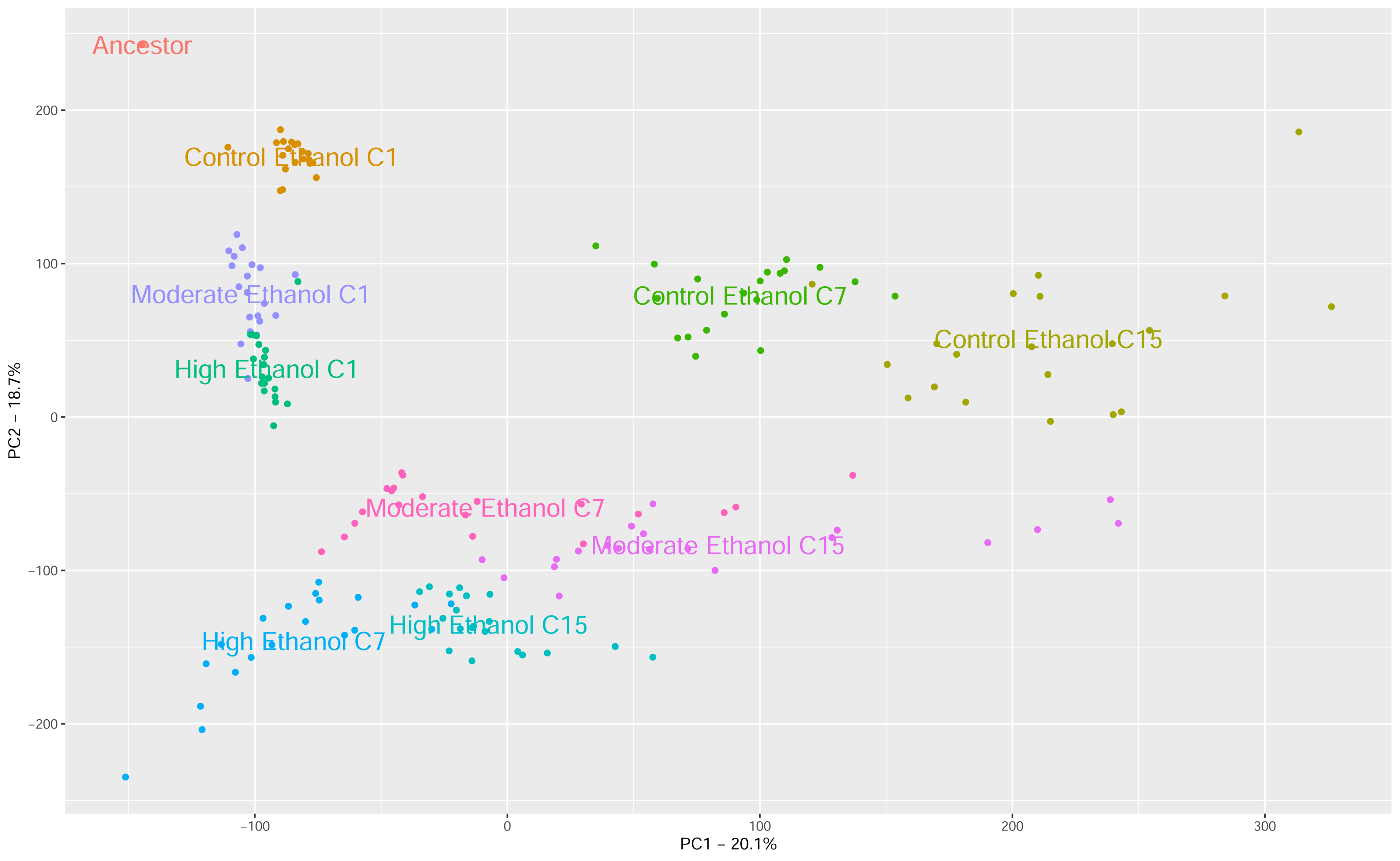
***PCA analysis of experimental populations***. Principal components analysis of raw allele frequencies (N=61,281) in the ancestor and replicate populations at each sequenced timepoint reveals distinct clustering by treatment. Early in the experiment (cycle 1=C1), populations cluster near to each other, and the ancestor, but by cycle 7 (C=7) and cycle 15 (C15) the replicates of each treatment cluster together.

The existence of treatment-specific responses supports our hypothesis that different selection intensities could lead to different targets of selection despite the same core stress. As shown in Figure 2, treatment-specific responses can be observed in each group. For instance, there are regions of the genome showing significant responses to selection in the H replicates (Figure 2A) and not the M replicates (Figure 2B) and vice versa despite both being exposed to ethanol stress. We also find cases where significant regions are found in the C replicates (Figure 2C) but not in either the H or M replicates despite all three treatments adapting to the same general laboratory conditions and maintenance protocols. Of note is the fact these control specific responses are actually our most significant treatment-specific responses.

An important caveat to consider is that interpreting our results as supportive of our hypothesis assumes that all of our response categories are biologically meaningful; however, this might not necessarily be the case. For instance, some SNPs in the high ethanol-specific category may in fact be better characterized as general ethanol alleles that are too weakly selected in M populations to be detected by our statistical tests. To account for this possibility, we calculated selection coefficient (*s*) estimates for each set of populations and compared how values varied between treatments and categories (see below).

### Selection coefficient estimates and response categories

To assess whether or not response categories are biologically meaningful, we first generated *s* estimates across the genome for the C, M and H populations across all polymorphic sites in the dataset (Supplementary Figure 6). Next, we calculated Pearson correlation coefficients (*r*) between *s* estimates from the C, M, and H populations across the genome and for the most significant SNPs for each major response category (Figure 4, see Supplementary Figure 7 for associated scatterplots). We define the most significant SNPs per category as those falling within 5KB of the most significant SNP per peak based on the CMH results (see Supplementary Table 3 for SNP counts and confidence intervals associated with *r* values). This was done in an effort to focus our analysis on sites likely to be most tightly linked to the true targets of selection.

**Figure 4.**
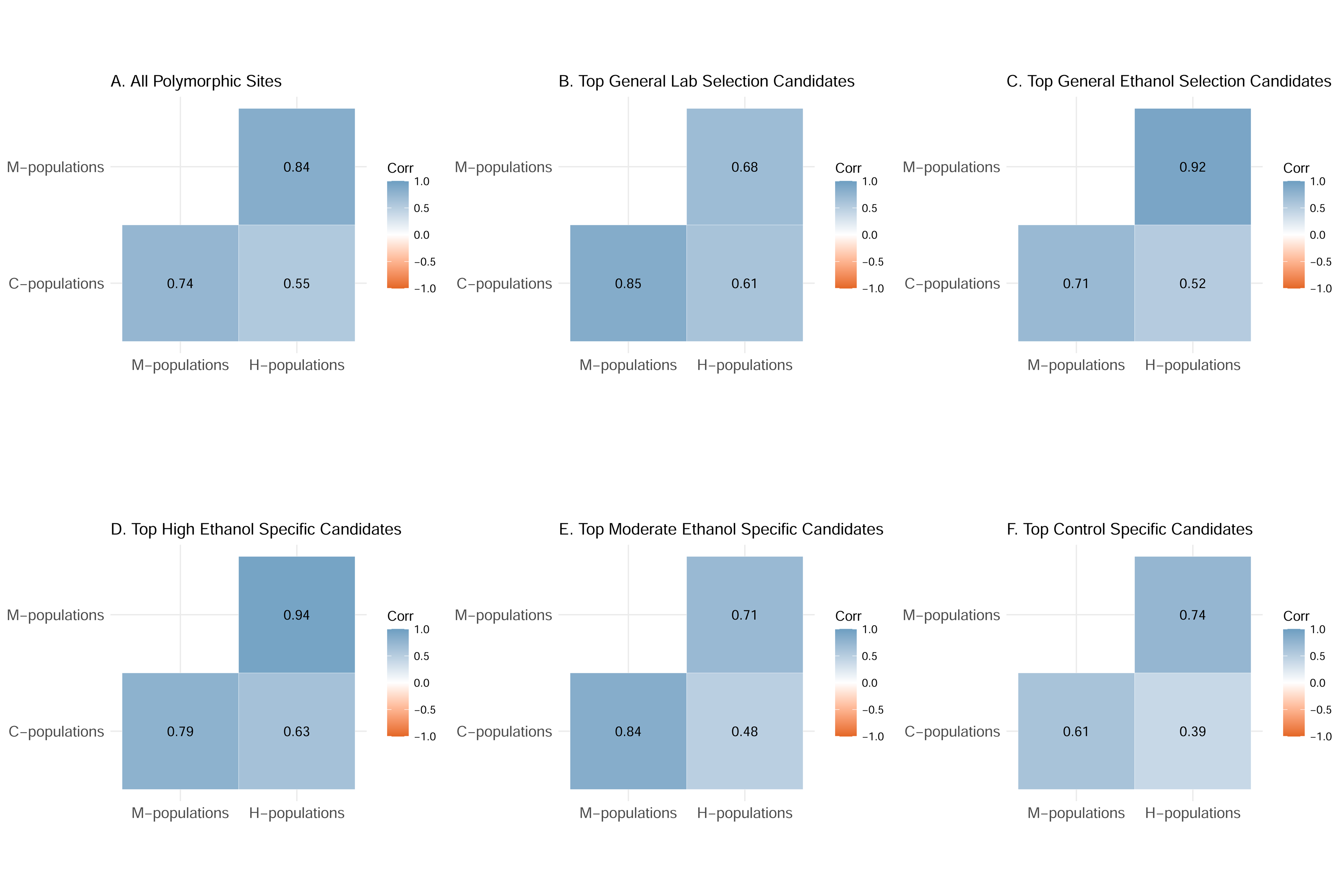
***Correlations between selection coefficients across categories***. We generated Pearson correlation coefficients (r) between selection coefficients estimated in C, M, and H populations for (A) all polymorphic sites, top (B) general selection candidates and (C) ethanol selection candidates, and top (D) high ethanol, (E) moderate ethanol, and (F) control specific candidates. Top candidate SNPs are defined as those in a 5 KB window around the most significant site for peaks associated with each category.

However, patterns are broadly the same if we perform the analysis using all SNPs in each category (Supplementary Figure 8).

The results of this analysis call into question the biological meaning of the response categories derived from our CMH and GLMM results. For instance, using the whole genome comparisons as a point of reference (Figure 4A, Supplementary Figure 7A), we find *s* estimates for the high ethanol-specific category are more highly correlated between the H and M populations than H or M vs C comparisons (Figure 4D). This, combined with values in the H population being generally higher (Supplementary Figure 7D) suggests that these sites may be under selection in both categories, but too weakly selected in the M population to be detected by our statistical tests. We find a similar pattern in the moderate ethanol-specific category where correlations indicate that candidate sites may also be under weaker selection in control conditions (Figure 4E, Supplementary Figure 7E). However, the signal is not as clear compared to the high ethanol-specific category. While these findings complicate the interpretation of our previously-defined response categories, they still indicate that selection intensity can shape outcomes through context-dependent effects on how strongly specific alleles are favored.

Perhaps the most surprising outcome of analyzing selection coefficient correlations in this way (Figure 4, Supplementary Figures 7-8) is that it reveals lower *s* estimates in the H populations than C and M, and lower correlations when comparing C or M populations to H than to each other. We interpret this as evidence that candidate alleles specifically beneficial in control conditions lose value as ethanol exposure increases (Figure 4F, Supplementary Figure 7F), and that this phenomenon drives most of the change observed in the experiment.

### Selection intensity and magnitude of allele frequency

To address our hypothesis that more intense selection should lead to greater changes in allele frequencies, we compared the mean change in SNP frequencies between the C, M, and H populations for our five major candidate SNP categories (Figure 5). Note, when performing statistical comparisons, we combine general ethanol stress and high ethanol-specific candidate SNPs into a single category. This was done as a result of our selection coefficient analysis suggesting that the latter are likely also responding in the M population, just to a lesser degree (see above). Based on Kruskal-Wallis tests, we see significant differences between treatments in all categories (p-values shown on Figure 5). So, we primarily relied on results from Wilcoxon tests between treatment pairs to assess whether or not greater selection intensity leads to greater changes in SNP frequencies at and around target sites.

**Figure 5.**
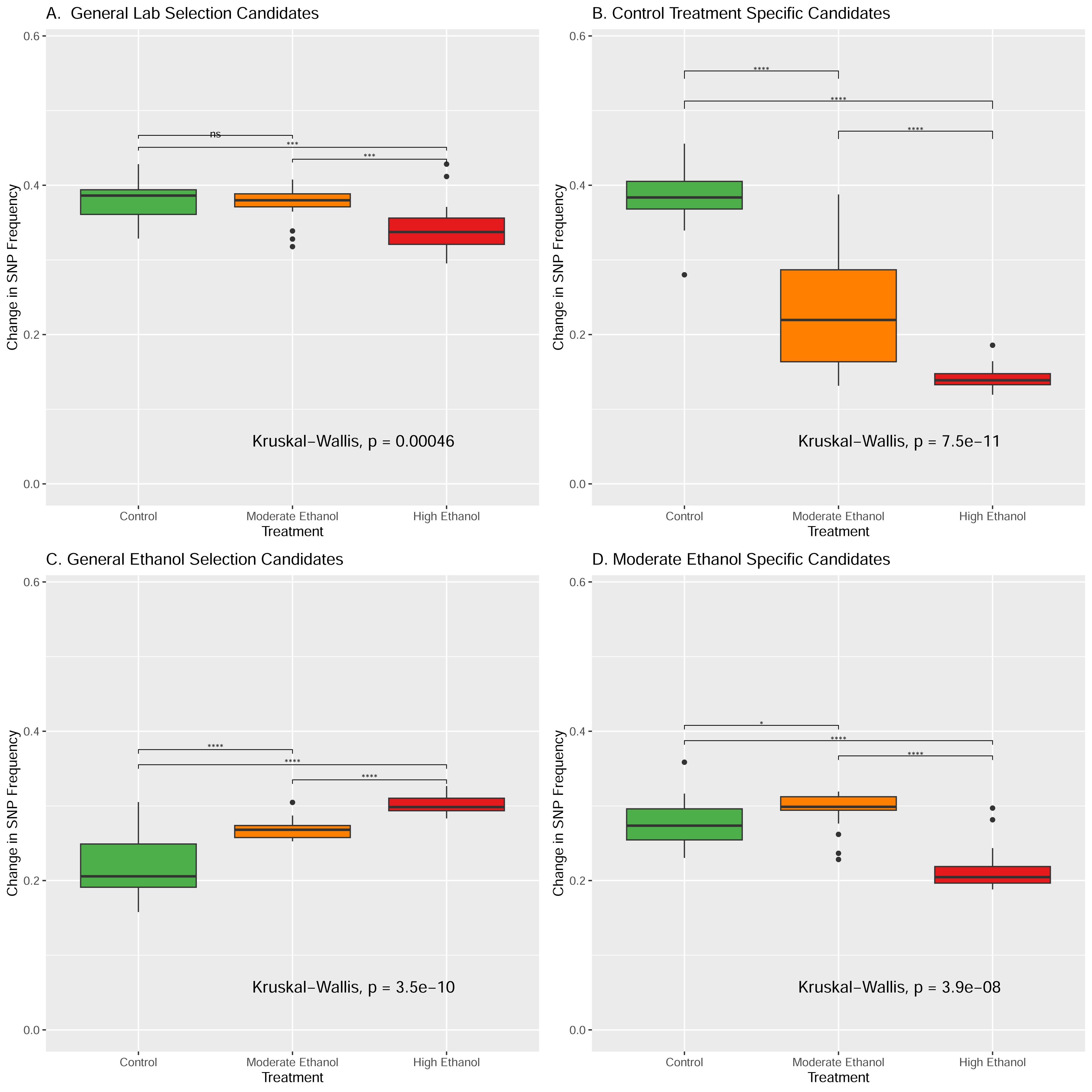
***Magnitude of allele frequency change across response categories.*** Boxplots show the mean absolute change in SNP frequency for each replicate in each treatment for (A) general laboratory adaptation candidates, (B) control-specific candidates, (C) general ethanol adaptation candidates, and (D) moderate ethanol-specific candidates. For this figure and analysis, we combined general ethanol and high ethanol-specific candidates into one group based on the high correlation between high and moderate-ethanol specific candidates revealed by Fig 4D.

With respect to the general laboratory selection category (Figure 5A), we find no significant difference in SNP frequency change between C and M populations (*p* = 0.52), but greater frequency changes in both of these treatments compared to the H populations (*p* = 0.008 and *p* = 5x10^-4^, respectively). As such, it appears that as ethanol exposure increases, these SNPs become less likely to respond. This pattern is further amplified in the control-specific category (Figure 5B) where we now find significant differences between C and M (*p* = 2.3x10^-8^), with a clear gradient in changes from C > M > H (*p* = 1.5x10^-11^ for C vs M comparison and *p* =2.6x10^-7^ for H vs C and H vs M comparisons).

As expected, for combined general ethanol selection and high ethanol-specific candidate SNPs (Figure 5C), we find greater changes in the H and M populations compared to C (*p* = 7.6x10^-7^ and *p* =3.9x10^-9^, respectively). We also see more change in the H populations than M (*p* = 9.9x10^-9^), indicating that that selection intensity impacts the magnitude of change. However, this does not extend to the moderate ethanol-specific candidates (Figure 5D). As with the *s* estimate comparisons, we again find some evidence that these are perhaps sites weakly selected for in both C and M populations. There is no significant difference in the level of change between C and M populations (*p* = 0.024), but both experience greater change than the H populations (*p* = 2.16x10^-7^ and *p* =9.8x10^-9^ for H vs C and H vs M comparisons). Lastly, it is perhaps worth noting that changes in frequency in general laboratory selection candidates are greater than changes in the general ethanol stress candidates. Like our CMH results (Figure 2), this further reinforces the idea that continued adaptation to laboratory conditions is a major driving force in this system.

### Candidate genes associated with different response categories

To compare potential differences in genes and pathways associated with our major response types (Table 1), we performed gene enrichment analyses using the candidate genes associated with each category. While we know that candidate SNPs in the high ethanol stress-specific category are likely also under selection in the M populations, we still analyzed this category separately as the results could potentially illuminate why these alleles respond less in the M populations. As shown in Supplementary Table 2, we find enrichment terms like “signaling” and “growth” that are shared across categories. However, we also find distinct enriched terms within response categories (both broadly and narrowly defined). This lends some creditability to the idea that selection is targeting different biological mechanisms across treatments. For instance, considering terms enriched in the general ethanol selection category, we find “alcohol catabolic process” and “response to organic substance” – terms that intuitively might underlie adaptation to long-term ethanol stress – that are not enriched in the general laboratory selection category. We also find distinct enrichment patterns with respect to treatment-specific categories. The most enriched terms in the high ethanol stress-specific response category are associated with cytokinesis and asexual reproduction, while the most enriched terms in the moderate ethanol stress-specific response category are more associated with pseudohyphal and filamentous growth, as well as metabolic function (e.g. lipid metabolism and sugar homeostasis). The most enriched terms in the control-specific response category are related to autophagy and immune function.

## Discussion

Using ethanol stress as the focal selective pressure, our experiment was designed to test if stronger selection was associated with greater changes in allele frequencies at target sites, and if targets of selection varied among treatments experiencing different selection intensities. However, our genomic analysis makes it clear that despite the use of an ancestral population that was previously subjected to a significant period of laboratory domestication (involving a dozen cycles of outcrossing and hundreds of asexual generations), adaptation to laboratory conditions is a major feature of our results (Figure 2 and Supplementary Figure 4C). While this complicates our interpretations, we believe our study still has the power to address our core questions, as the control treatment allows us to distinguish between adaptive responses associated with laboratory domestication and those associated with ethanol stress (Table 1). And while we cannot definitively say ethanol exposure is exerting a greater selective pressure than general laboratory handling, it stands to reason that total selection intensity is highest in the high ethanol stress treatment, followed by the moderate ethanol stress treatment, and lowest in the control treatment; in other words, we have created a spectrum of stressful environments, in which some elements are shared, and others are unique.

### Patterns of growth in evolved populations imply different avenues of adaptation

The observed growth-related phenotypes in our experimentally-evolved populations suggest complex evolutionary dynamics underlying adaptation to ethanol stress. In YPD (the control medium), replicates of the C, M, and H treatments all have similar doubling times after 15 cycles and all grow slightly faster than the ancestral population (Figure 1). This is perhaps expected given that our genomic results indicate all populations are continuing to adapt to laboratory conditions. However, under moderate and high ethanol stress conditions, patterns are more complicated. Under moderate ethanol stress, replicates of the H treatment have slower growth rates and lower carrying capacities (Figure 1C-D, and Supplementary Figure 3B) than all other groups assayed. And while they do have faster doubling times than controls and the ancestral population in high ethanol stress conditions (Figure 1E-5), they again have the lowest carrying capacities (Supplementary Figure 3C). Replicates of the M treatment have similar doubling times under high ethanol conditions, but do not display the same level of reduction in carrying capacity (Supplementary Figure 3C). Therefore, we conclude that adaptation to ethanol stress involves mechanisms not directly related to growth rate. Differences between the M and H populations across media types also implicate potential trade-offs between growth rate and final sustainable population size. Notably, the relationship between carrying capacity (as defined by our analysis) and fitness is not known; this metric simply reflects the value at which growth appears to cease in the population. The simplest interpretation of carrying capacity is that it reflects the point at which nutrients have been depleted from the medium, preventing further growth; therefore it is reasonable to expect that the fastest growing populations might also have the lowest carrying capacities, though this is not always what we observe. We think there could be many possible mechanisms underlying differentiation in carrying capacity, including changes in cell size, shape, structure and/or aggregation – and any of these may or may not trade-off with growth rate.

These observations of potential trade-offs support the idea that pleiotropy and genetic interactions may underlie adaptation to these distinct environments. This is consistent with other E&R work that explores stress resistance; for example, Kawecki et al. (2021) find that even though the same genes underlie starvation resistance in experimentally-evolved *D. melanogaster*, directionality of allele frequency shifts change depending on whether selection is imposed at larval or adult life stages due to underlying trade-offs. More broadly, we do not find that increased selection intensity results in a clear and proportional phenotypic response. As such, selection intensity should be carefully considered when using experimental evolution to predict how specific traits might change in response to a given selective pressure.

### The magnitude of allele frequency change at candidate sites is context-dependent

If increased selection intensity does result in greater changes in SNP frequency at and around targets of selection as simulation work by Christodoulaki et al. (2019) suggests, we would expect that at general ethanol candidate sites, changes should be greater in the high ethanol stress treatment compared to the moderate ethanol treatment. In principle, this should also be true for candidate sites associated with general laboratory adaptation; we would predict that changes in the high and moderate ethanol stress treatments should be greater than those observed in the control treatment. Based on our selection coefficient analysis, we combined general ethanol and high ethanol-specific candidate sites. Comparing change in SNP frequencies among these candidates, we find the expected gradient where H populations have the greatest change and C the smallest (Figure 5C). As such, here we find clear support for the idea that greater selection intensity leads to greater change at and around target sites. However, results beyond this suggest more complicated dynamics.

For the general laboratory selection category, we see a reversal of our expected pattern with the smallest changes in the H populations and largest in the C populations (Figure 5A). This may reflect some level of antagonism between adaptation to lab conditions and ethanol stress, and this antagonism increases with higher levels of ethanol exposure. We also observe this pattern in control-specific candidates (Figure 5B), with even smaller changes in the M and H populations compared to the general laboratory selection candidates. Moderate ethanol-specific candidates also support this general narrative as frequency changes are significantly lower in the H populations compared to C and M (which do not significantly differ from one another); this could indicate sites weakly selected under lab conditions and low levels of ethanol exposure, but not adaptative at high ethanol levels. This outcome aligns with results from Christodoulaki et al. (2019), who simulated allele frequency changes associated with different distances from new trait optima and found small shifts at target sites when trait optima are close. Altogether, we interpret these results as evidence that different selection intensities can potentiate distinct, though perhaps often subtle, adaptive responses.

We also considered the possibility that in this experiment, adaptation to laboratory protocols may have been stressful enough that the inclusion of ethanol exposure had little additional impact (i.e. all treatments are under very intense selection). This could explain why sites in the general laboratory selection and control-specific categories show the greatest changes in SNP frequency on average. However, growth rate assays make it clear that the ancestral populations and controls are indeed negatively impacted by ethanol exposure (Figure 1), and PCA analysis shows that populations in ethanol treatments cluster distinctly from those in the control treatments. As such, we generally argue that the three experimental treatments experienced distinct and specific selection pressures, and that our results partially support the idea of a positive correlation between selection intensity and allele frequency change.

On the issue of shared adaptation to laboratory conditions, it is worth noting that such responses are not readily observed in similar work from Linder et al. (2022). Here the authors, also working with outcrossing yeast, characterize adaptation to different chemical stress treatments. Unlike our study where we find many shared responses to selection across treatments, they at most only observe parallel responses across 4 to 5 treatments. We believe there are several possible explanations for this difference.

First, perhaps they were simply more successful in adapting their ancestral population to laboratory conditions before deriving experimental populations. In both their study and ours, ancestral populations were maintained in the lab for similar numbers of generations before experiments were started, so differences might be due to the specifics of maintenance protocols (e.g. Linder et al. use a liquid handling robot and small volumes while we carried out manual transfers). Additionally, while we characterized responses to selection to different concentrations of ethanol, they studied responses to 17 entirely different chemical stressors. The absence of shared responses across all treatments could bedue to a lower likelihood of a given “lab adaptation” allele being beneficial in so many different environments. This seems especially plausible given our observed control-specific responses to selection. We also note that their experiment features very high selection intensities by design, with dosages that populations can barely survive in early generations. As a direct point of comparison, our high ethanol stress treatment involves 10% ethanol while theirs is 12.5%. Finally, as their haplotype-based analysis implicates that selection impacts very large swaths of the genome in each treatment, it is difficult to identify specific targets of selection and completely rule out the possibility of common laboratory adaptation.

### The presence of different selection response categories reinforces that trait architectures vary with selection intensity

We interpret the existence of treatment-specific responses to selection across groups as evidence that the genetic architecture of adaptation varies with selection intensity. For instance, while we do find evidence of general ethanol adaptation, we also find candidates that appear specific to either high or moderate ethanol exposure (red peaks in Figure 2A, orange peaks in Figure 2B). The observed correlation in *s* estimates between treatments for these candidates (Figure 4D-E and Supplementary Figure 8D-E), leads us to conclude that high ethanol-specific candidates are also under weaker selection in the M populations, and likewise many moderate ethanol-specific candidates are also under weaker selection in the controls. This calls into question the biological meaning of our categories, perhaps highlighting our limited ability to identify weakly-selected sites with our statistical methods. However, even with this caveat, this observed context-dependence supports the idea that selection intensity, broadly defined, shapes adaptive outcomes. The apparent phenotypic trade-offs observed in the M and H treatments reinforces this interpretation.

Surprisingly, the clearest evidence for treatment-specific adaptive response comes from our control populations (Figure 2C). Unlike the moderate or high ethanol-specific peaks which are often small or contain SNPs from multiple response categories, control-specific responses include some of our most significant and “pure” peaks. So, while these regions of the genome are strongly associated with the maintenance and laboratory conditions that are shared by all experimental treatments, they appear to be (at most) weakly selected under moderate ethanol stress and not selected at all under high ethanol stress (see Figure 4F and Supplementary Figure 7F for evidence from *s* estimates). Here one possibility is while these alleles do confer advantages under control conditions, they have costs in the presence of ethanol and those costs increase as the level of ethanol exposure increases (i.e. context-dependence due to pleiotropy- associated trade-offs). Alternatively, this pattern could also be explained by negative epistatic interactions that amplify with increasing levels of ethanol stress. While we do not have the ability to distinguish between these two scenarios, both support the idea that selection intensities can create distinct environments favoring distinct genetic architectures of adaptative traits, and concomitant GxE interactions.

### Selection targets different biological mechanisms across treatments

As shown by our gene enrichment analysis, different biological functions appear to be associated with adaptation across treatments. Focusing on the M and H populations, we find that adaptation to general ethanol stress involves genes related to alcohol catabolic processes, signaling, and responses to abiotic stimuli (Supplementary Table 4). However, we also observe distinct enrichment terms within the M and H categories; while adaptation to high ethanol stress is associated with GO terms relating to the integrity of cell division (e.g. cytokinesis, DNA repair, and budding), these functions may either be unnecessary or come with some cost at lower levels of ethanol stress. Similarly, while adaptation to moderate ethanol stress is associated with GO terms related to metabolic function, these are not implicated at high levels of ethanol exposure. Together, these results provide further evidence that the genetic architecture of adaptation varies with selection intensity due to underlying differences and potential trade-offs in the mechanisms through which adaptation occurs.

With respect to the various mechanisms associated with ethanol resistance in and across the moderate and high ethanol stress treatments, our findings are consistent with past work suggesting ethanol resistance in *S. cerevisiae* is complex and involves many genetic and physiological mechanisms (reviewed in Ding et al. 2009 and Ma and Liu 2010). Signaling, lipid metabolism, carbohydrate homeostasis all stand out as mechanisms implicated by past studies. Work focusing on adaptation to high ethanol exposure in asexual *S. cerevisiae*, in this case peaking at 12% of the culture medium, also specifically implicated mutations in genetic pathways associated with cell cycle and DNA replication (Voordeckers et al. 2015). Here the authors suggest that delayed cell cycle progression allows for greater protection for individual cells. This sort of survival mechanism, which might more accurately be described as stress tolerance rather than stress resistance, may drive the apparent lower carrying capacity observed in the populations adapted for high ethanol stress in the presence of ethanol. This stands out as an intuitive example of a potential context-dependent adaptative mechanism associated with ethanol stress.

## Conclusion

Our results generally support the idea that selection intensity significantly impacts evolutionary dynamics. As such, this should be a major point of consideration when synthesizing results across E&R studies even when they feature the same stress or condition as a focal selective pressure. It should also be considered when attempting to extrapolate findings from E&R studies to natural populations. For instance, candidate sites and dynamics observed in E&R studies with intense selection are perhaps unlikely to translate very well to a natural population facing more moderate environmental changes and vice versa. Our observations also indicate a greater need for models of polygenic adaptation that include the possibility of extensive pleiotropy and epistasis. Furthermore, while here we chose to focus on selection intensity, it is likely that other dynamic circumstances of natural environments, such as temporal heterogeneity in selection, will also affect outcomes in meaningful ways. As such, we believe our findings illuminate a clear need for studies that explicitly demonstrate how experimental and population-genetic factors shape evolutionary dynamics in E&R studies moving forward. Lastly, as with the work of Pfenninger and Foucault (2020), our results demonstrate the importance of appropriate controls even when timeseries data are available. Our controls ultimately prevented us from making a number of spurious associations between large portions of observed genomic responses to selection and ethanol stress resistance.

## Data Availability

The raw sequence files generated over the course of this project are available through NCBI SRA (BioProject ID: PRJNA839395) and scripts used to process raw data and perform SNP calling are available through Github (https://github.com/mollyburke/Burke-Lab-SNP-calling-pipeline). Core scripts necessary to reproduce our results are available through Github (https://github.com/mphillips67/Genomics-Ethanol-Stress-Outcrossing-Yeast) and major input and results files are available through Dryad (https://doi.org/10.5061/dryad.tqjq2bw27).

## Author Contributions

M.A.P. and M.K.B. conceived of this project. M.A.P., R.K.B., M.S., S.Z. and M.S.P. did the lab work necessary to generate the data sets used in this study. M.A.P. carried out the data analysis, and M.A.P. and M.K.B. wrote the manuscript.

### Acknowledgements

We thank Oregon State University’s Center for Quantitative Life Sciences for use of their computational and sequencing resources. This work was supported by startup funds provided to M.K.B. by the College of Science at Oregon State University, as well as by National Institutes of Health award R35GM147402. M.A.P. was supported by a National Science Foundation Postdoctoral Fellowship (NSF 1906246).

## Supplementary Figures

**Supplementary Figure 1.**
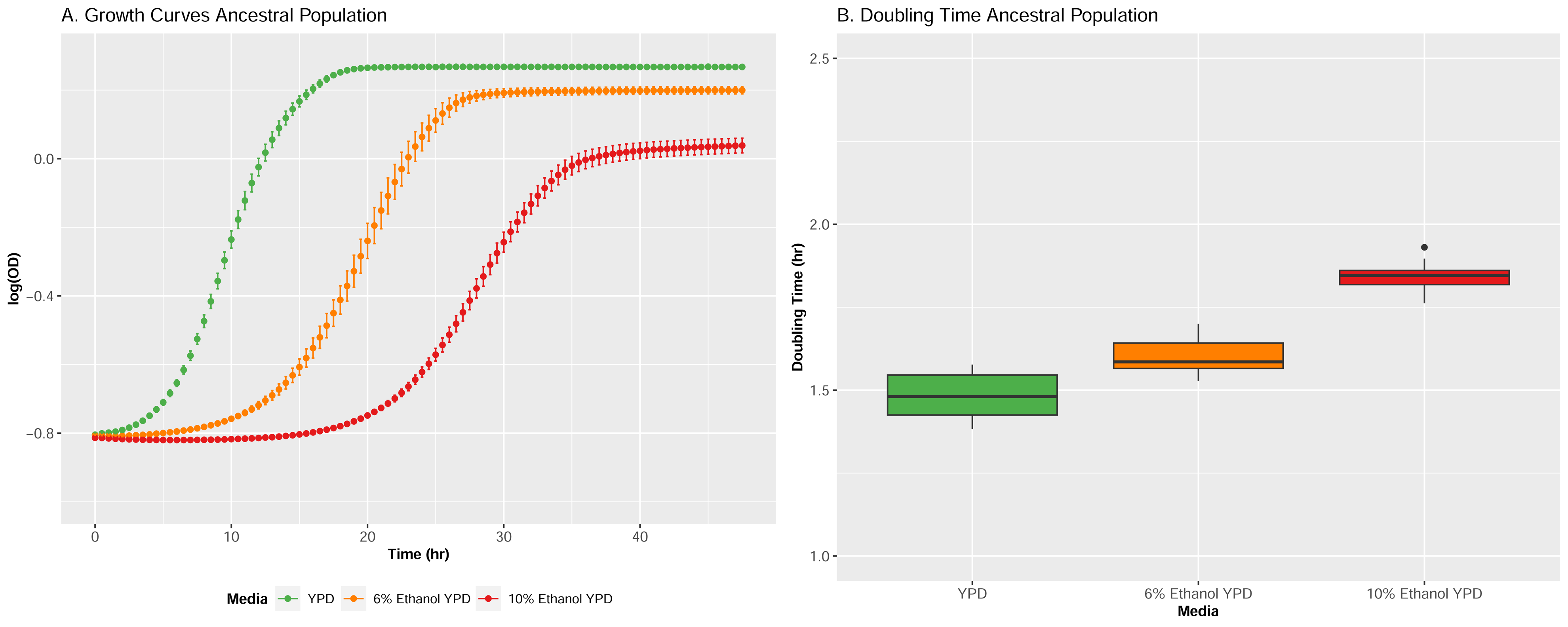
Growth rates (A) and doubling times (B) for the ancestral population in plain YPD, 6% ethanol YPD, and 10% ethanol YPD (note: this figure is based on assays performed before the start of the evolution experiment; different data were used to produce Figure 1 and Supplementary Figure. 3).

**Supplementary Figure 2.**
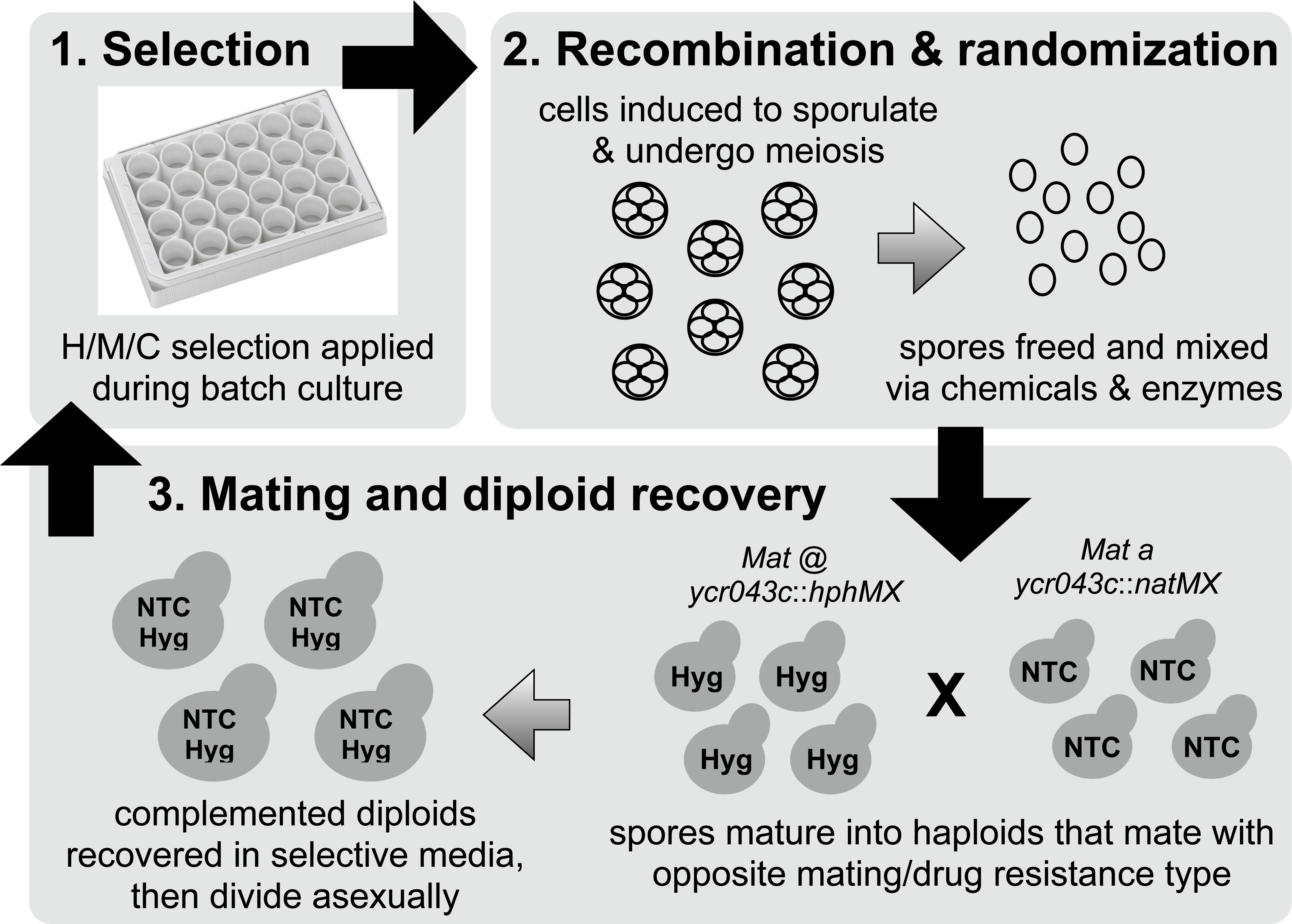
General schematic showing how the H, M, and C populations were maintained over the course of this study.

**Supplementary Figure 3.**
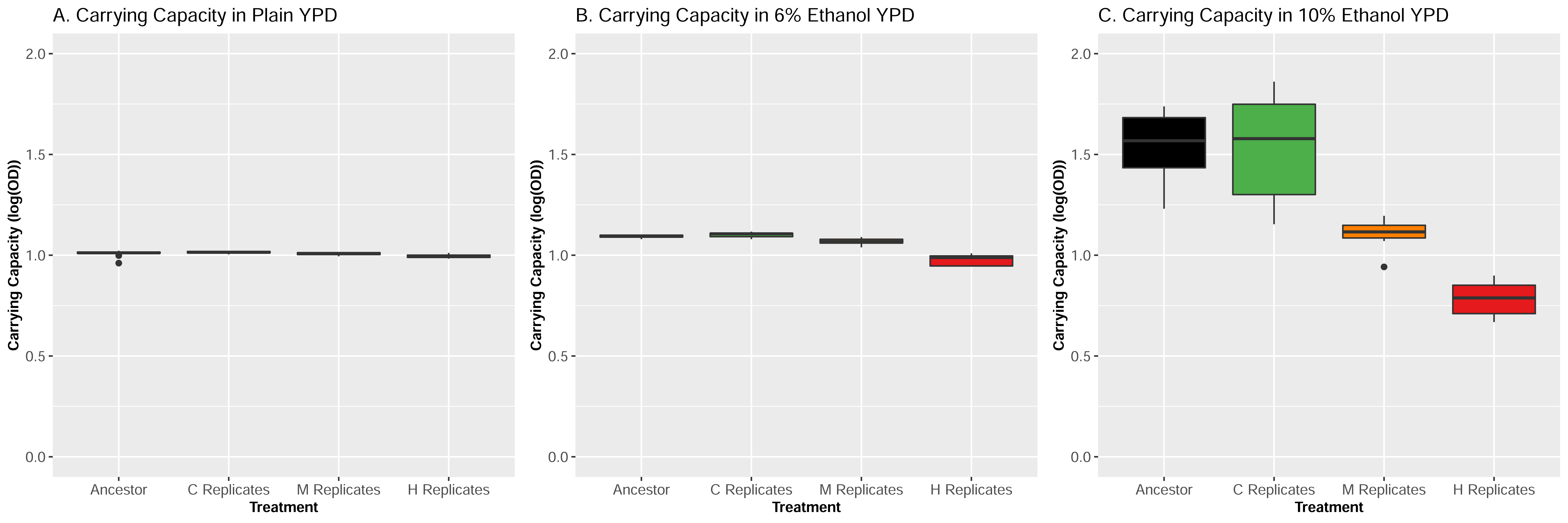
Carrying capacity estimates based on three independent growth rate assays for the ancestral population and experimental populations after 15 cycles of selection in control (A), moderate ethanol (B), and high ethanol (C) conditions. In each assay, nine technical replicates of the ancestor and 10 randomly-chosen replicates from each treatment were evaluated.

**Supplementary Figure 4.**
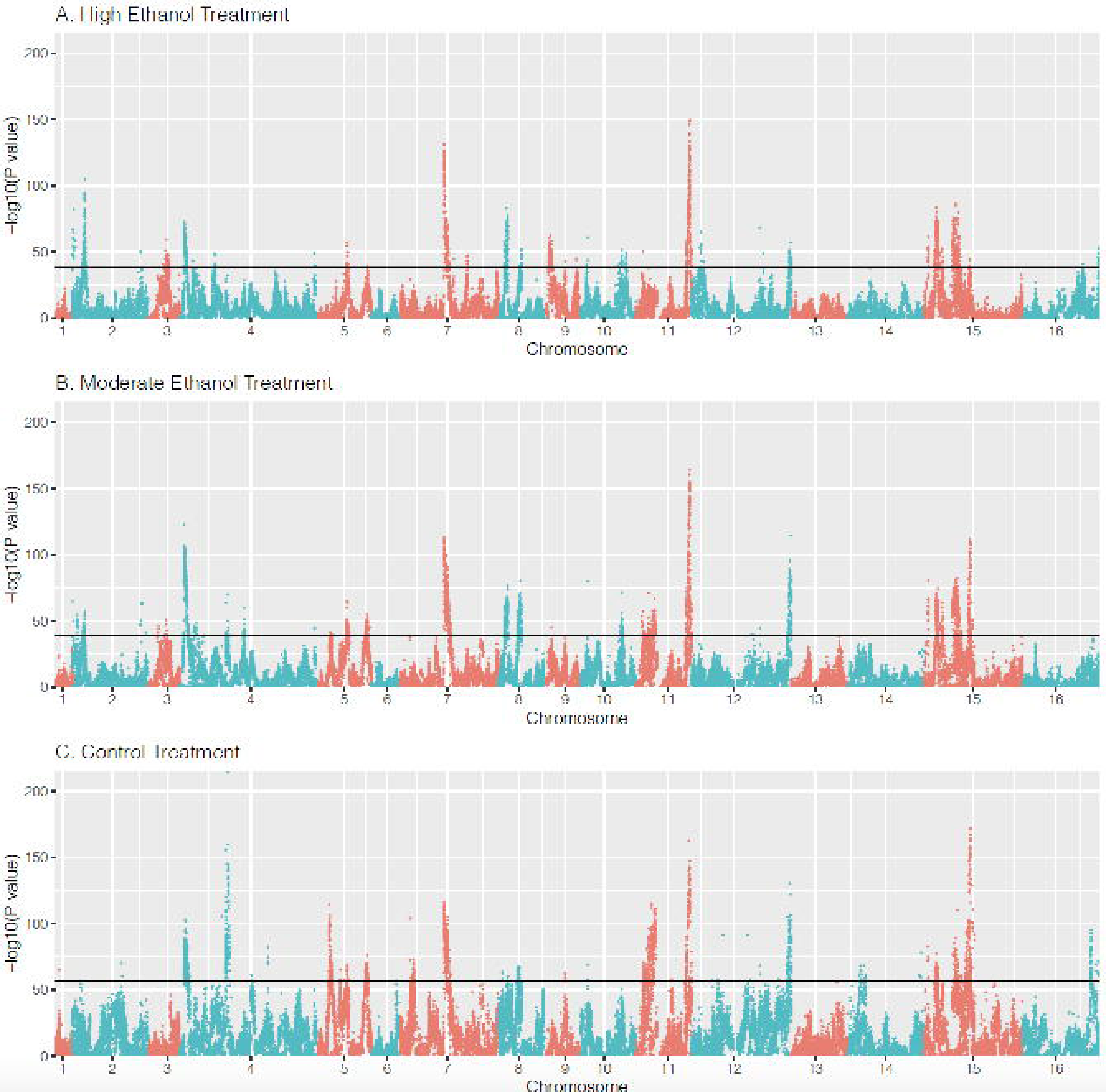
CMH results comparing SNP frequencies between cycles 1 and 15 in (A) High ethanol stress, (B) Moderate ethanol stress, and (C) Control treatments. Black lines represent permutation derived significance thresholds.

**Supplementary Figure 5.**
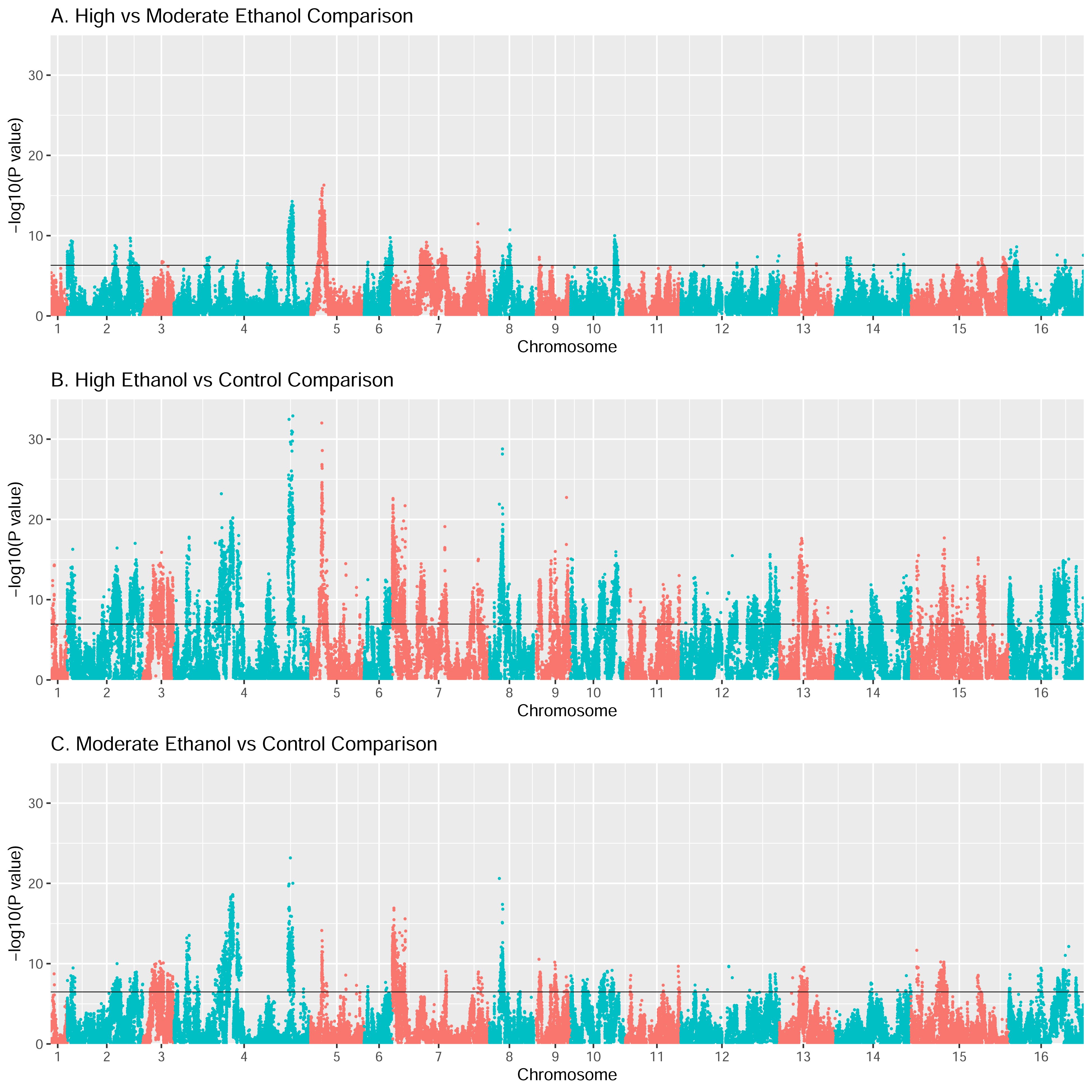
GLMM results from comparing SNP frequencies cycle 15 SNP frequencies between (A) High and Moderate Ethanol Stress treatments, (B) High Ethanol Stress and Control treatments, and (C) Moderate Ethanol Stress and Control treatments. Black lines represent permutation derived significance thresholds.

**Supplementary Figure 6.**
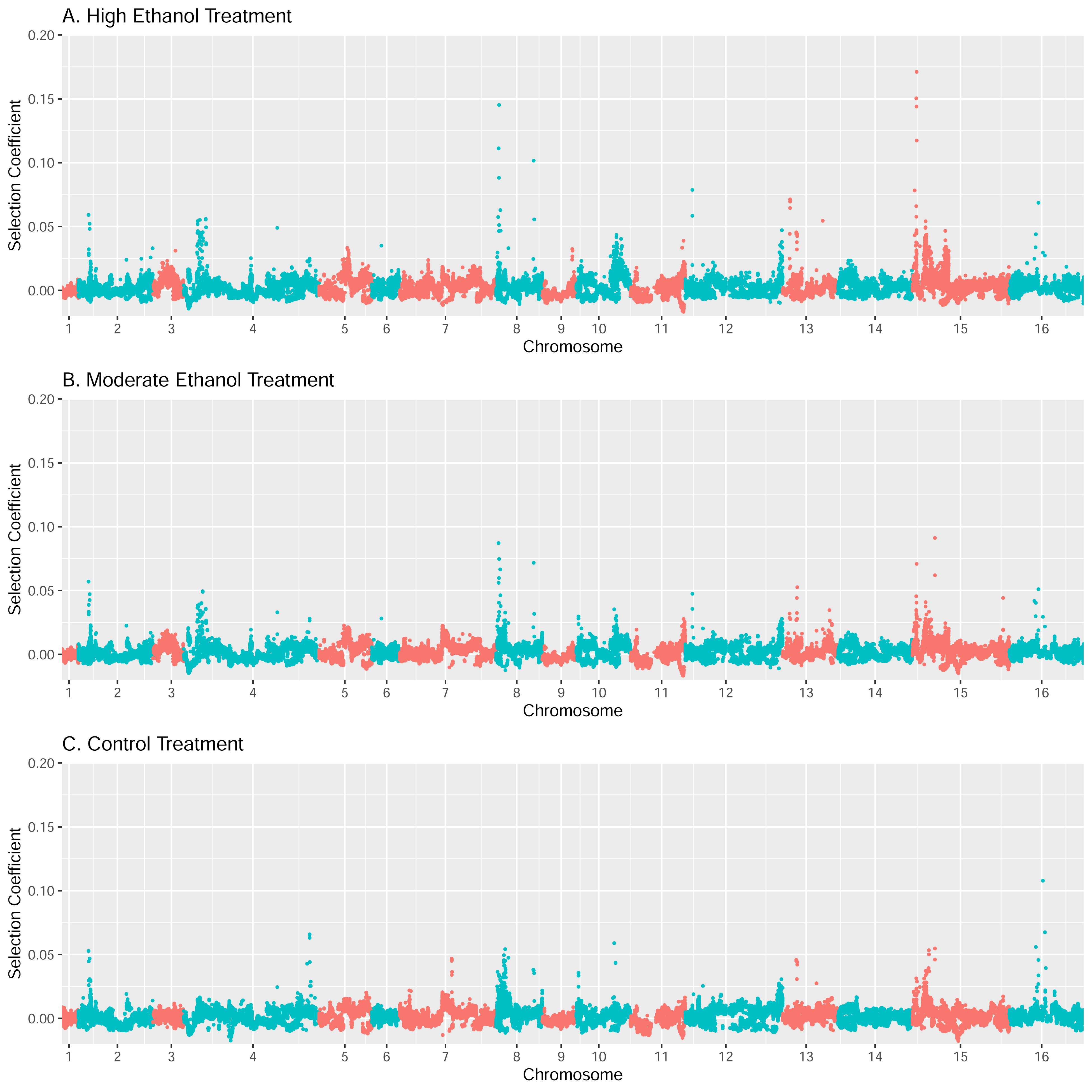
Selection coefficient estimates across the genome based on changes in SNP frequencies between cycles 1 and 15 in (A) high ethanol, (B) moderate ethanol, and (C) control treatments.

**Supplementary Figure 7.**
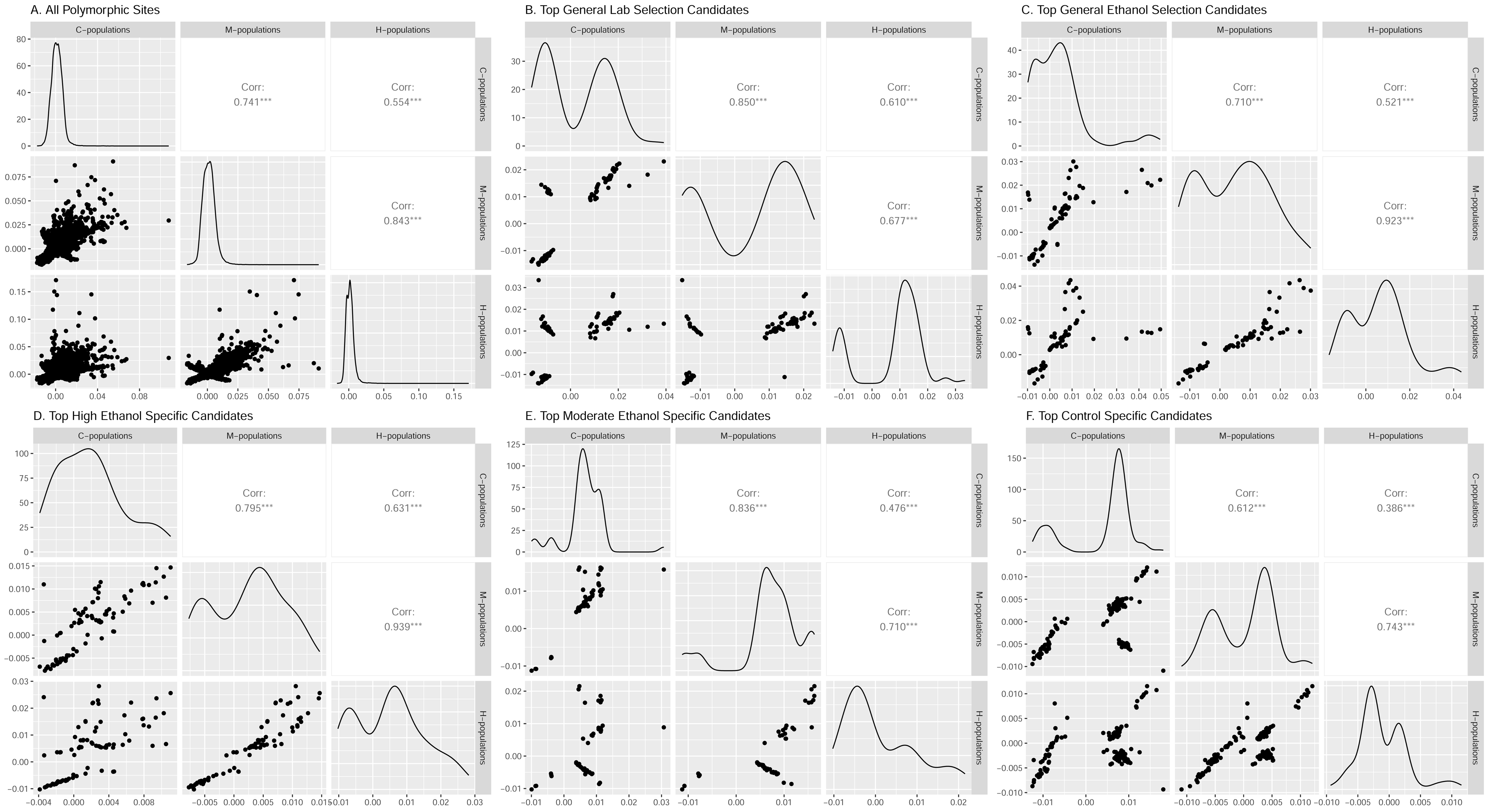
Scatter plots comparing groups, density plots, and correlations based on selection coefficient estimates for C, M, and H populations for (A) all polymorphic sites, top (B) general selection candidates and top (C) ethanol selection candidates, and (D) high ethanol, (E) moderate ethanol, and (F) control specific candidates. Top candidate SNPs are defined as those in a 5 KB window around the most significant site for peaks associated with each category.

**Supplementary Figure 8.**
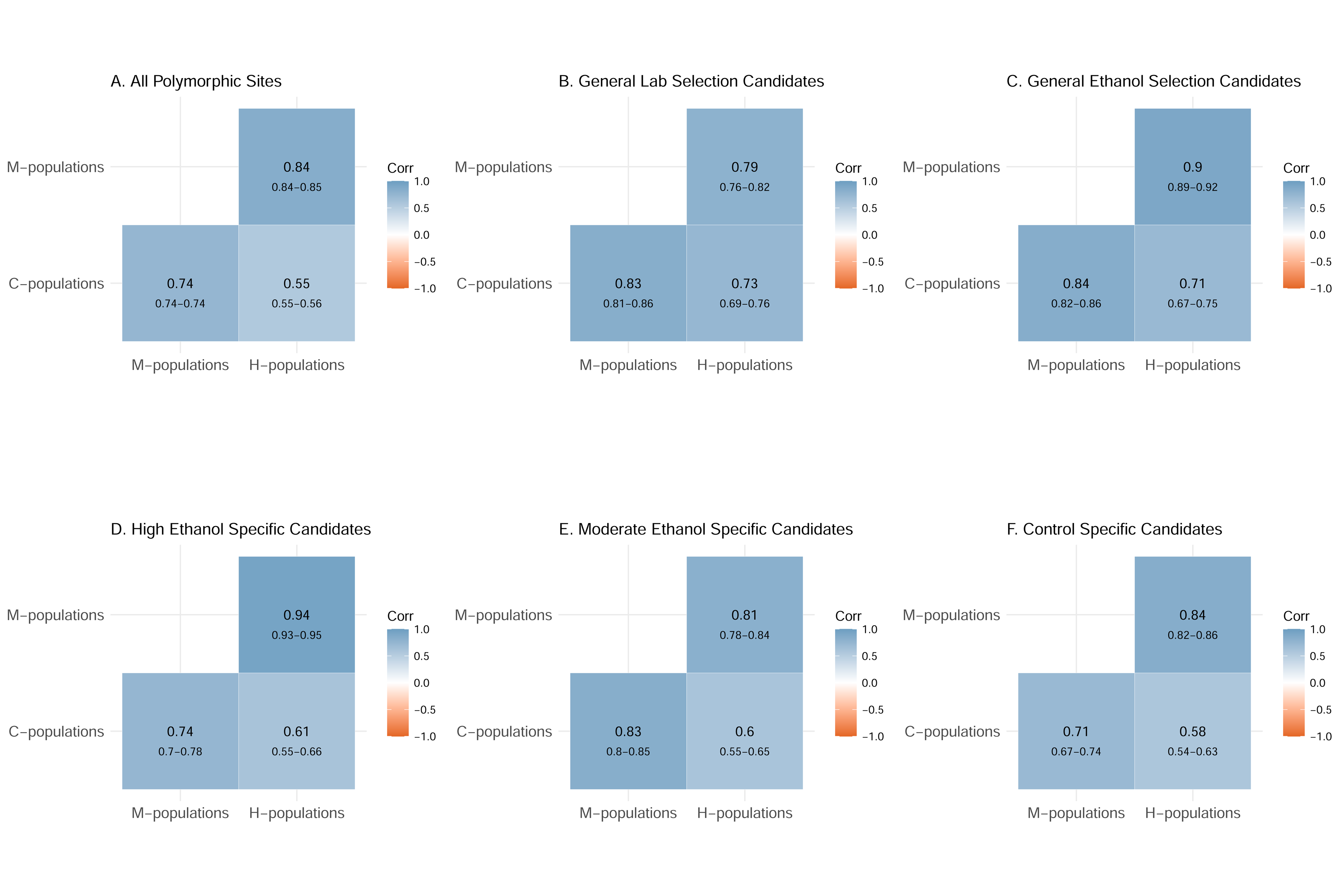
Correlations between selection coefficient estimates for C, M, and H populations for (A) all polymorphic sites, (B) general selection candidates and (C) ethanol selection candidates, and (D) high ethanol, (E) moderate ethanol, and (F) control-specific candidates. 95% confidence intervals associated with each value are also displayed.

## References

1. Barghi, N., Hermisson, J., Schlötterer, C. (2020). Polygenic adaptation: a unifying framework to understand positive selection.

2. Barata, C., Borges, R., & Kosiol C. (2023) Bait-ER: a Bayesian method to detect targets of slection in evolve-and-sequence experiments. Journal of Evolutionary Biology, 36,29–44.

3. Bates, D., Machler, M., Bolker, B., & Walker, S. (2015). Fitting linear mixed-effects models using lme4. Journal of Statistical Software, 67, 1–48.

4. Baym, M., Kryazhimskiy, S., Liberman, T.D., Chung, H., Desai, M.M., & Kishony, R. (2015). Inexpensive multiplexed library preparation for megabase-sizes genomes. PLoS One, 10, e0131262.

5. Benjamin, J., Berger, J.O, Johannesson, M., Nosek, B.A., Wagenmakers, E.J., … Johnson, W.E. (2018) Redefined statistical significance. Nature Human Behavior 2, 6–10.

6. Boyle, E.A., Li, Y.I., & Pritchard, J.K. (2017) An expanded view of complex traits: from polygenic to omnigenic. Cell, 169, 1177–1186 .

7. Burke, M.K. (2012), How does adaptation sweep through the genome? Insights from long-term selection experiments. Proceedings of the Royal Society B, 279, 5029–5038.

8. Burke, M.K., Liti, G., & Long, A.D. (2014). Long standing genetic variation drives repeatable experimental evolution in outcrossing populations of *Saccharomyces cerevisiae*. Molecular Biology and Evolution, 32, 3228–3239.

9. Burke, M.K., McHugh, K.M., & Kutch, I.C. (2020). Heat shock improves random spore analysis in diverse strains of Saccharomyces cerevisiae. Frontiers in Genetics, 11, 1636.

10. Burke, M.K. (2023). Embracing complexity: Yeast evolution experiments featuring standing genetic variation. Journal of Molecular Evolution 91:281–292

11. Christodoulaki, E., Barghi, N, & Schlötterer, C (2019). Distance to trait optimum is a crucial factor determining the genomic signature of polygenic adaptation. bioRxiv, 10.1101/721340.

12. Cingolani, P., Platts, A., Wang, L., Coon, M., Nguyen, T., Wang, L., Land, S.J., Ruden, D.M., & Lu, X. (2012). A program for annotating and predicting the effects of single nucleotide polymorphisms, SnpEff: SNPs in the genome of *Drosophila melanogaster* strain w1118; iso-2, iso-3. Fly, 6, 80–92.

13. Cubillos, F.A., Louis, E.J., & Liti, G. (2009) Generation of a large set of genetically tractable haploid and diploid *Saccharomyces* strains. FEMS Yeast Research, 9, 1217–1225.

14. Ding, J., Huang, X., Zhang, L., Zhao, N, Yang, D., & Zhang, K. (2009). Tolerance and stress response to ethanol in the yeast Saccharomyces cerevisiae. Applied Microbiology and Biotechnology, 85, 253–263.

15. Graves, J.L., Hertweck, K.L., Phillips, M.A., Han, M.V., Cabral, L.G., Barter, T.T., … Rose MR. 2017. Genomics of parallel experimental evolution in Drosophila. Molecular Biology and Evolution, 34, 831–842.

16. Hayward, L.K., & Sella, G. (2021). Polygenic adaptation after a sudden change in environment. bioRxiv, 10.1101/792952.

17. Jha, A.R., Miles, C.M., Lippert, N.R., Brown, C.D., White, KP, & Kretiman, M. (2015). Whole-genome resequencing of experimental populations reveals polygenic basis of egg-size variation in *Drosophila melanogaster*. Molecular Biology and Evolution, 32, 2616–2632.

18. Kawecki, T.J., Erkosar, B., Dupuis, C., Hollis, B., Stillwell, R.C., Kapun, M. (2021). The genomic architecture of adaptation to larval malnutrition points to a trade-off with adulat starvation resistance in *Drosophila*. Molecular Biology and Evolution, 28, 2732–2749.

19. Kessner, D., & Novembre, J. (2015). Power analysis of artificial selection experiments using efficient whole genome simulation of quantitative traits. Genetics, 199, 991–1005.

20. Linder, RA., Majumder A., Chakraborty, M., & Long A.D. (2020). Two synthetic 18-way outcrossed populations of diploid budding yeast with utlity for complex trait dissection. Genetics 215, 323–342.

21. Linder, R.A., Zabanavar, B., Majumder, A., Shyan-Chiao Hoang, H., Delgado, V., Tran, R., La, V., Leemans, S., & Long, A.D. (2022). Adaptation in outbred sexual yeast is repeatable, polygenic and favors rare haplotypes. Molecular Biology and Evolution, 39, msac248.

22. Liu, H., Maclean, C.J., Zhang, J. (2019). Evolution of the yeast recombination landscape. Molecular Biology and Evolution 36(2):412–422

23. Long AD, Liti G, Luptak A, Tenaillon O. (2015). Elucidating the molecular architecture of adaptation via evolve and resequence experiments. Nature Reveiews Genetics, 16, 567–582.

24. Ma, M., & Liu, Z.L. (2010). Mechanism of ethanol tolerance in *Saccharomyces cerevisiae*. Applied Microbiology and Biotechnology, 87, 829–845.

25. Otte, KA., Nolte, V., Mallard, F., & Schlötterer, C. (2021). The genetic architecture of temperature adaptation is shaped by population ancestry and not by selection regime. Genome Biology 22, 211.

26. Pfenninger, M. &, Foucault, Q. (2020). Genomic processes underlying rapid adaptation of a natural *Chironomus riparius* population to unintendedly applied experimental selection pressures. Molecular Ecology, 29, 536–548.

27. Phillips, M.A., & Burke, M.K. (2021). Can laboratory evolution experiments teach us about natural populations? Molecular Ecology, 30, 877–879.

28. Phillips, M.A., Kutch, I.C., Long, A.D., & Burke, M.K. (2020) Increased time sampling in an eolve-and-resequence experiment with outcrossing *Saccharomyces cerevisiae* reveals multiple paths of adaptive change. Molecular Ecology, 29, 4898–4912.

29. Phillips, M.A., Kutch, I.C., McHugh, K.M., Taggard, S.K., & Burke, M.K. (2021). Crossing design shapes patterns of genetic variation in synthetic recombinant populations of *Saccharomyces cerevisiae*. Scientific Reports, 11, 19551.

30. Phillips, M.A., Briar, R.K., Scaffo, M., Zhout, S. & Burke, M.B. (2022). Strength of selection potentiates distinct adaptive responses in an evolution experiment with outcrossing yeast. NCBI SRA, BioProject ID: PRJNA839395.

31. R Core Team. (2021). R: a language and environment for statistical computing. R foundation for Statistical Computing, Vienna, Austria. www.R-project.org.

32. Schlötterer, C., Kofler, R., Versace, E., Tobler, R., & Franssen, S.U. (2015). Combining experimental evolution with next-generation sequencing: a powerful tool to study adaptation from standing genetic variation. Heredity, 114, 431–440.

33. Schlötterer, C. (2023). How predictable is adaptation from standing genetic variation? Experemtnal evolution in Drosohila highlights the central role of redundnacy and linkage disequilibrium. Philosophical Transactions of the Royal Society B, 378, 20220046.

34. Sprouffske, K. & Wagner, A. (2016). Growthcurver: an R package for obtaining interpretable meterics from microbial growth curves. BMC Bioinformatics, 17, 172 (2016).

35. Stetter, M.G., Thornton, K., & Ross-Ibarra, J. (2018). Genetic architecture and selective sweeps after polygenic adaptation to distant trait optima. PLoS Genetics, 14, e1007794.

36. Taus, T., Futschik, A., & Schlötterer, C. (2017). Quantifying selection with pool-seq time series data. Molecular Biology and Evolution, 34, 3023–3034.

37. Van der Auwera, GA., & O’Connor, B.D. (2020). Genomics in the cloud: using Docker, GATK, WDL, and Terra. O’reilly Media.

38. Visscher, P.M., & Yang, J. (2016) A plethora of pleiotropy across complex traits. Nature Genetics, 48, 707–708.

39. Vlachos, C., & Kofler, R. (2019). Optimizng the power to identify the genetic basis of complex traits with evolve and resequence studies. Molecular Biology and Evolution, 36, 2890–2905.

40. Vlachos, C., Burny, C., Pelizzola, M., Borges, R., Futschik, A., Kofler, R., & Schlötterer, C. (2019). Benchmarking software tools for detecting and quantifying selection in evolve and resequecing studies. Genome Biology, 20, 169.

41. Voordeckers, K., Komine, J., Das, A., Espinosa-Cantu, A., De Maeyer, D., Arslan, A., Van Pee, M., van der Zande, E, Meert, W., Yang, Y., Zhu, B., Marchal, K., DeLuna, A., Verstrepen, K.J. (2015). Adaptation to high ethanol reveals complex evolutionary pathways. PloS Genetics, 11, e1005635.

42. Wiberg, R.A., Gaggiotti, O.E., Morrisssey, M.B., Ritchie M.G. (2017). Identifying consistent allele frequency differences in studies of stratified populations. Methods in Ecology and Evolution, 8, 1899–1909.

